# Profiling the initial burst of beneficial genetic diversity in clonal cell populations to anticipate evolution

**DOI:** 10.1101/2020.07.10.196832

**Authors:** Daniel E. Deatherage, Jeffrey E. Barrick

**Affiliations:** Department of Molecular Biosciences, Center for Systems and Synthetic Biology, The University of Texas at Austin, Austin, Texas 78712, U.S.A.

## Abstract

Clonal populations of cells continuously evolve new genetic diversity, but it takes a significant amount of time for the progeny of a single cell with a new beneficial mutation to outstrip both its ancestor and competitors to fully dominate a population. If these driver mutations can be discovered earlier—while they are still extremely rare—and profiled in large numbers, it may be possible to anticipate the future evolution of similar cell populations. For example, one could diagnose the likely course of incipient diseases, such as cancer and bacterial infections, and better judge which treatments will be effective, by tracking rare drug-resistant variants. To test this approach, we replayed the first 500 generations of a >70,000-generation *Escherichia coli* experiment and examined the trajectories of new mutations in eight genes known to be under positive selection in this environment in six populations. By employing a deep sequencing procedure using unique molecular identifiers and target enrichment we were able to track 301 beneficial mutations at frequencies as low as 0.01% and infer the fitness effects of 240 of these. Distinct molecular signatures of selection on protein structure and function were evident for the three genes in which beneficial mutations were most common (*nadR, pykF,* and *topA*). We detected mutations hundreds of generations before they became dominant and tracked beneficial alleles in genes that were not mutated in the long-term experiment until thousands of generations had passed. This type of targeted adaptome sequencing approach could function as an early warning system to inform interventions that aim to prevent undesirable evolution.

## Introduction

New genetic variation naturally arises in lineages of cells and organisms during genome replication and repair. These *de novo* mutations are the main drivers of adaptive evolution in many populations, particularly those with little or no recombination or standing genetic variation. In large laboratory populations of asexual microbes, numerous lineages with different beneficial mutations arise and contend within a population before any one outcompetes the ancestor and its competitors (Good et al., 2017; Lang et al., 2013; Maddamsetti et al., 2015). This ‘clonal interference’ leads to heterogeneous populations with many lineages simultaneously adapting via distinct sets of mutations, though often these mutations are in a small subset of genes that are under the strongest selection (Desai et al., 2012; Gerrish and Lenski, 1998; Park and Krug, 2007).

In human cancers and chronic microbial infections, single cells clonally expand in a similar fashion by evolving driver mutations that lead to disease progression and drug resistance. Both solid tumors and blood cancers have been shown to be genetically heterogeneous (Marusyk et al., 2012; Merlo et al., 2006; Thomas et al., 2006). *De novo* mutations taking over normal cell populations can lead to carcinogenesis (Genovese et al., 2014; Watson et al., 2020), and mutations in cancer cells drive neoplastic progression (Merlo et al., 2010), differences in responses to chemotherapy (Landau et al., 2013), and relapse (Ding et al., 2012). Populations of *Pseudomonas aeruginosa* and other bacteria that persistently infect the lungs of cystic fibrosis patients become increasingly invasive and antibiotic resistant over time (Marvig et al., 2015; Stefani et al., 2017; Winstanley et al., 2016). In these cases, there are also specific genetic loci that are repeatedly mutated in different individuals. Better predicting the future evolution of each of these types of cell populations and others would inform treatment decisions and improve medical outcomes.

Cells used in biomanufacturing are also prone to evolving unwanted genetic heterogeneity (Renda et al., 2014; Rugbjerg and Sommer, 2019). Typically, these cells have been heavily engineered to optimize the titer of a product of interest at the expense of rapid cellular replication (Lee and Kim, 2015; Nielsen and Keasling, 2016). Therefore, there are strong selective pressures for ‘escape mutations’ that cause production to decline. Usually escape mutations directly inactivate one or more key genes in the engineered pathway. The resulting nonproducing cells can become dominant during the many cell divisions that are necessary to scale these processes up to large bioreactors (Rugbjerg et al., 2018; Sandoval et al., 2014; Zelder and Hauer, 2000). The ability to predict the future evolution of nonproducing cells before attempting scale-up could guide strain design decisions and thereby improve the efficiency of industrial processes.

Evolution experiments conducted in laboratory environments reproduce key aspects of microbial evolution that are observed in chronic infections and bioreactors (Barrick and Lenski, 2013; Gresham and Dunham, 2014). Certain aspects of genomic and phenotypic evolution in these systems are surprisingly predictable while others are not (Barrick, 2020; Cvijović et al., 2018; Furusawa et al., 2018; McDonald, 2019; Rainey et al., 2017). In theory, profiling many rare mutations in the earliest stages of clonal interference using high-throughput DNA sequencing should allow one to anticipate the future evolution of a population and other populations that evolve in similar environments. Existing studies have generally been limited to reliably identifying mutations that are present in at least one sample at a frequency above ~1-10% (due to sequencing depth costs and per-base error rates) when they have already succeeded in becoming dominant (Barrick and Lenski, 2009; Chubiz et al., 2012; Good et al., 2017; Lang et al., 2013; Traverse et al., 2013). Theory and simulations predict that many more highly beneficial mutations evolve in these populations but never reach such high frequencies before they are driven extinct (Desai et al., 2012; Gerrish and Lenski, 1998), and recent studies that track the evolution of barcoded lineages of microbes show that this is the case (Levy et al., 2015; Venkataram et al., 2016).

Here, we used methods for selectively increasing sequencing depth and lowering sequencing error rates to deeply profile the initial burst of rare beneficial mutations in laboratory populations of *E. coli.* We directly identified diverse beneficial mutations in eight genes when they were orders of magnitude lower in frequency and hundreds of generations earlier than could be accomplished by standard metagenomic sequencing methods. Molecular signatures in the large sets of beneficial mutations found in three of these genes made it possible to understand the nature of selection acting on their functions in this environment. By comparing our results to the long history of a >70,000-generation *E. coli* evolution experiment that used the same ancestral strains and nearly identical culture conditions (Lenski et al., 1991), we are able to evaluate the potential of this type of targeted adaptome analysis for anticipating the evolution of cell populations.

## Results

### Replaying the beginning of a long-term evolution experiment

We initially examined the evolution of nine replicate *E. coli* populations that were propagated via daily serial transfers in glucose-limited minimal medium for 500 generations. Our experiment used the same *E. coli* strains as the Lenski long-term evolution experiment (LTEE) and similar growth conditions (see **Methods**). Each population was inoculated with a 50/50 mixture of the two neutrally marked LTEE ancestor strains to visualize the initial selective sweep (Hegreness et al., 2006). Most populations maintained a roughly equal representation of descendants of both ancestral strains through the first 300 generations (**Fig. 1**). These dynamics are in agreement with what has previously been observed in studies of the LTEE, where few mutations reach a high frequency in the first few hundred generations of evolution (Good et al., 2017). We did not further analyze three populations. Two of these (A4 and A5) were purposefully omitted because they exhibited early sweeps of one marker type, which indicates that their dynamics might have been dominated by one or a few “jackpot” mutations that occurred very early during outgrowth of these populations from single cells. The third omitted population (A8) exhibited typical marker dynamics.

**Figure 1.**
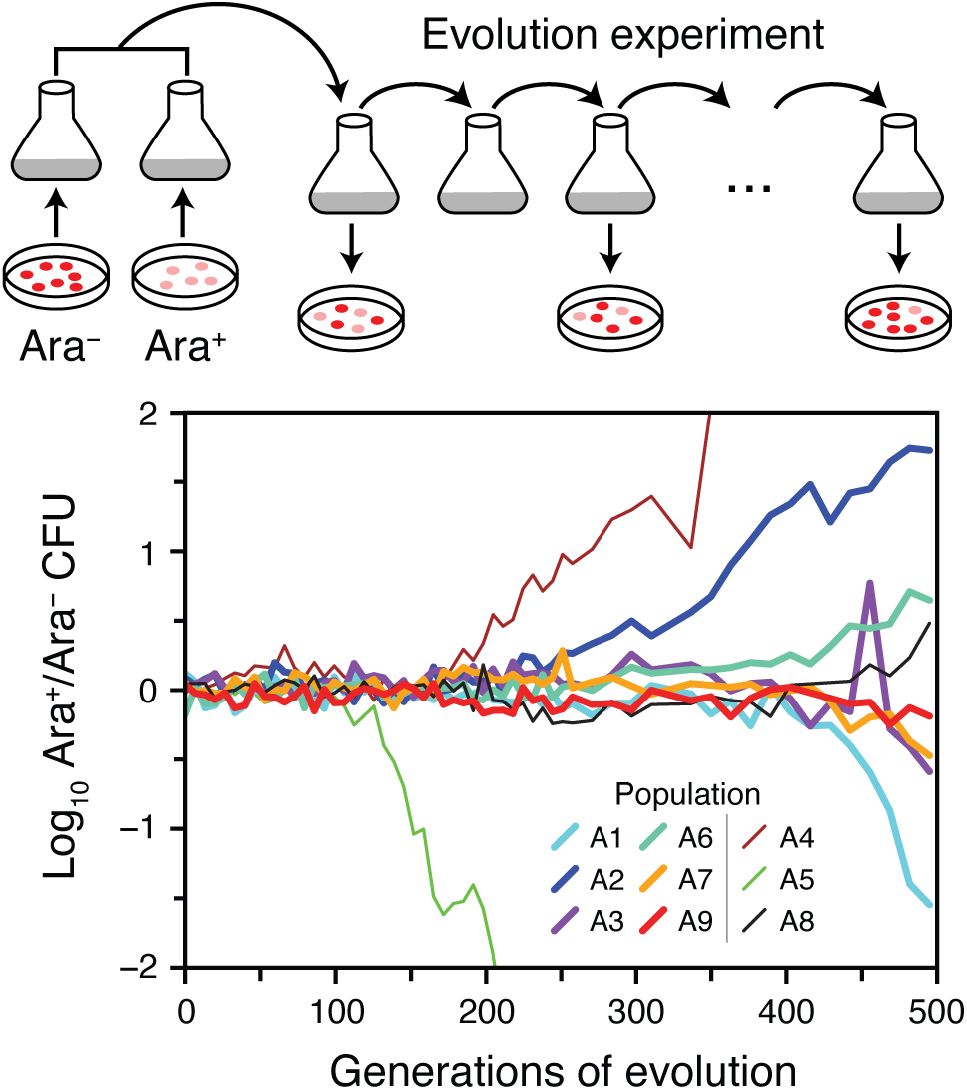
Replaying the first selective sweep of a long-term evolution experiment. Nine *E. coli* populations were initiated from equal mixtures of two variants of the ancestral strain that differ in a neutral genetic marker for arabinose utilization (Ara). We observed the evolutionary dynamics of these populations over ~500 generations of regrowth from 75 daily 1:100 serial transfers by periodically plating dilutions of each population on indicator agar. The ratio of Ara^+^ cells (pink colonies) to Ara^-^ cells (red colonies) diverges from 1:1 when descendants of one ancestor type accumulate enough of a fitness advantage due to *de novo* beneficial mutations that they take over. We focused further analysis on six of the nine populations (thick lines).

### Reconstructing the trajectories of new beneficial mutations

We next performed deep sequencing of eight genes at ~25 generation increments over the entire 500 generations of the evolution experiment for four of the six populations that we examined further. These eight genes (*nadR, pykF, topA, spoT, fabR, ybaL, hslU,* and *iclR*) are known to be targets of selection in the LTEE (Good et al., 2017; Tenaillon et al., 2016). Illumina libraries containing unique molecular identifiers (Schmitt et al., 2012) were prepared for sequencing and enriched for the regions of interest using solution based hybridization (Bainbridge et al., 2010). Consensus sequence reads were generated based on groups of reads with identical unique molecular identifiers and aligned to the *E. coli* genome to predict mutations, including using split-read mapping to identify transposon insertions and large deletions (**Fig. 2A**). The enrichment procedure was effective: an average of 73.5% of consensus reads per sample mapped to the targeted regions that together constitute only 0.780% of the 4.63 Mbp genome. In the sample with the median number of total consensus reads, the average coverage depth across each of the eight genes of interest exceeded 5,000 (**Fig. 2B**). After analyzing patterns in mutation frequencies over time to eliminate other systematic biases (see Methods), we were able to track the evolution and competition of 181 *de novo* mutations, including when many were present in less than 0.1% of the cells in a population (**Fig. 2C**, **Fig. 3**).

**Figure 2.**
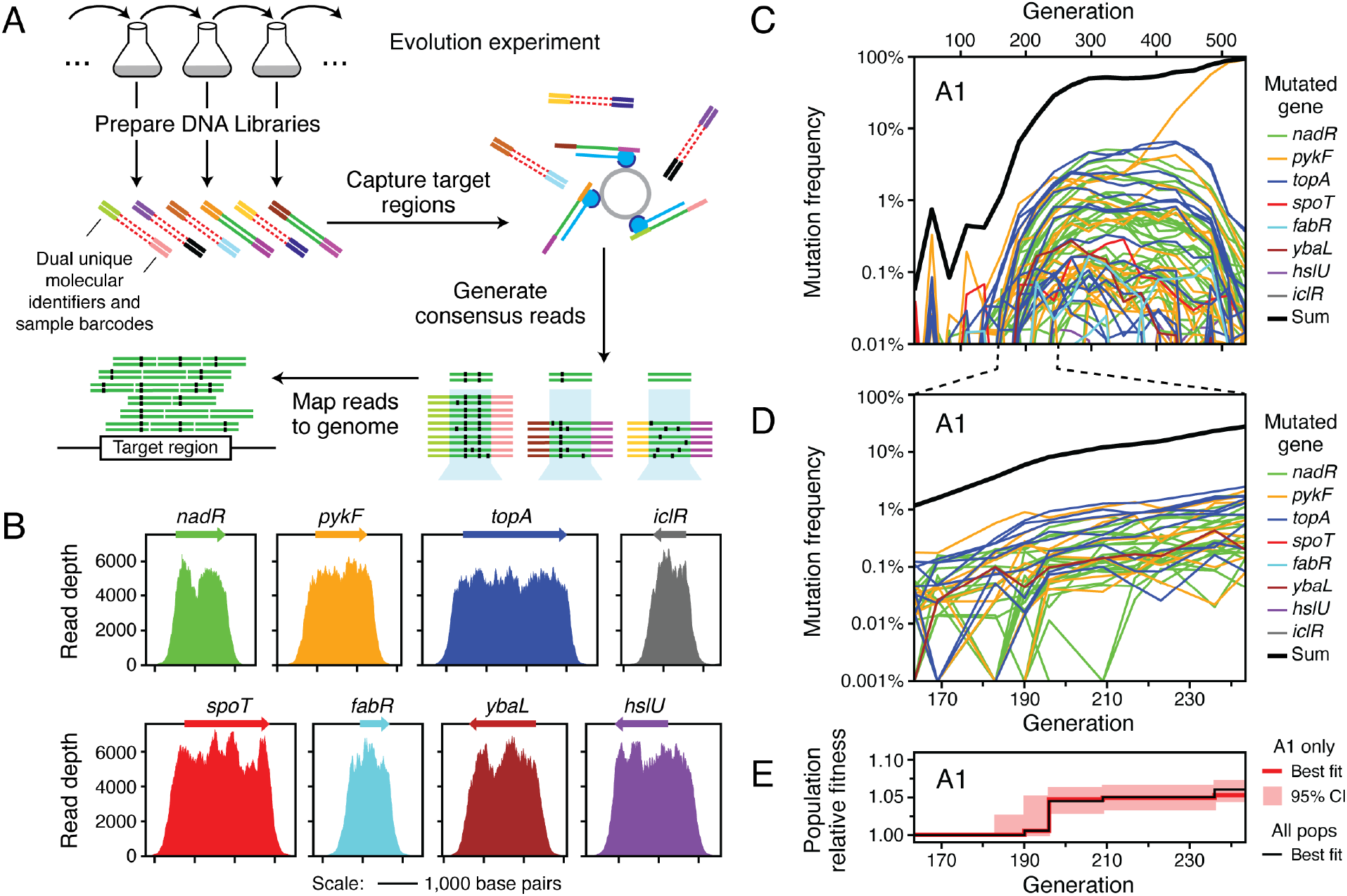
Profiling many beneficial mutations in the first selective sweep by deep sequencing. (A) Schematic of the deep sequencing approach. Genomic DNA is directly isolated from the *E. coli* populations and prepared for paired-end Illumina sequencing with sample barcodes and dual unique molecular identifiers (colored ends attached to red/green double stranded DNA). DNA fragments matching the targeted genome regions (green centers) are captured by probes (blue) bound to magnetic beads and other sequences are washed away (red centers). Reads in pairs that have the same dual unique molecular identifiers, which implies that they were PCR amplified during library preparation from the same original genomic DNA fragment, are used to construct consensus reads to eliminate sequencing errors. Consensus reads are mapped to the reference genome to call sequence variants. (B) Enrichment of reads mapping to eight genes known to be early targets of selection in this environment from the long-term evolution experiment. The final coverage depth of consensus reads in and around these genes is shown for a typical sample (population A7 at 500 generations). (C) Frequency trajectories for mutations in the eight targeted genes as well as the sum total frequency in population A1 over the complete time course of the evolution experiment. When a mutation was not detected at a time point, its trajectory is shown as passing through a frequency of 0.0001% (outside of the plot bounds). (D) Mutation frequency trajectories for population A1 during the window from 133 to 213 generations when mutations were first reaching detectable frequencies as they outcompeted the ancestral genotype. At time points when a mutation was not detected, its frequency is shown as 0.001% (at the bottom of the plot). (E) Estimated relative fitness of population A1 in each interval between the time points in the window time course. The frequency trajectories of all beneficial mutations in the initial sweep shown in D were used to jointly estimate the average fitness of the entire population from the deceleration in the rate of increase of the observed mutation trajectories as genotypes with beneficial mutations became common (see **Methods**). This fitness trajectory fit accounts for all cells in the population, regardless of whether they have a mutation in the targeted genes or elsewhere in the genome. The red line is the maximum likelihood estimate of the population fitness trajectory. The red shading around it shows 95% confidence intervals on this value in each interval. The black line shows the increase in fitness estimated for a consensus model that was jointly fit to all mutations tracked in all six populations for which sequencing data was collected in this window. The consensus population fitness trajectory was used when estimating the fitness effects of individual mutations (see **Methods**)

**Figure 3.**
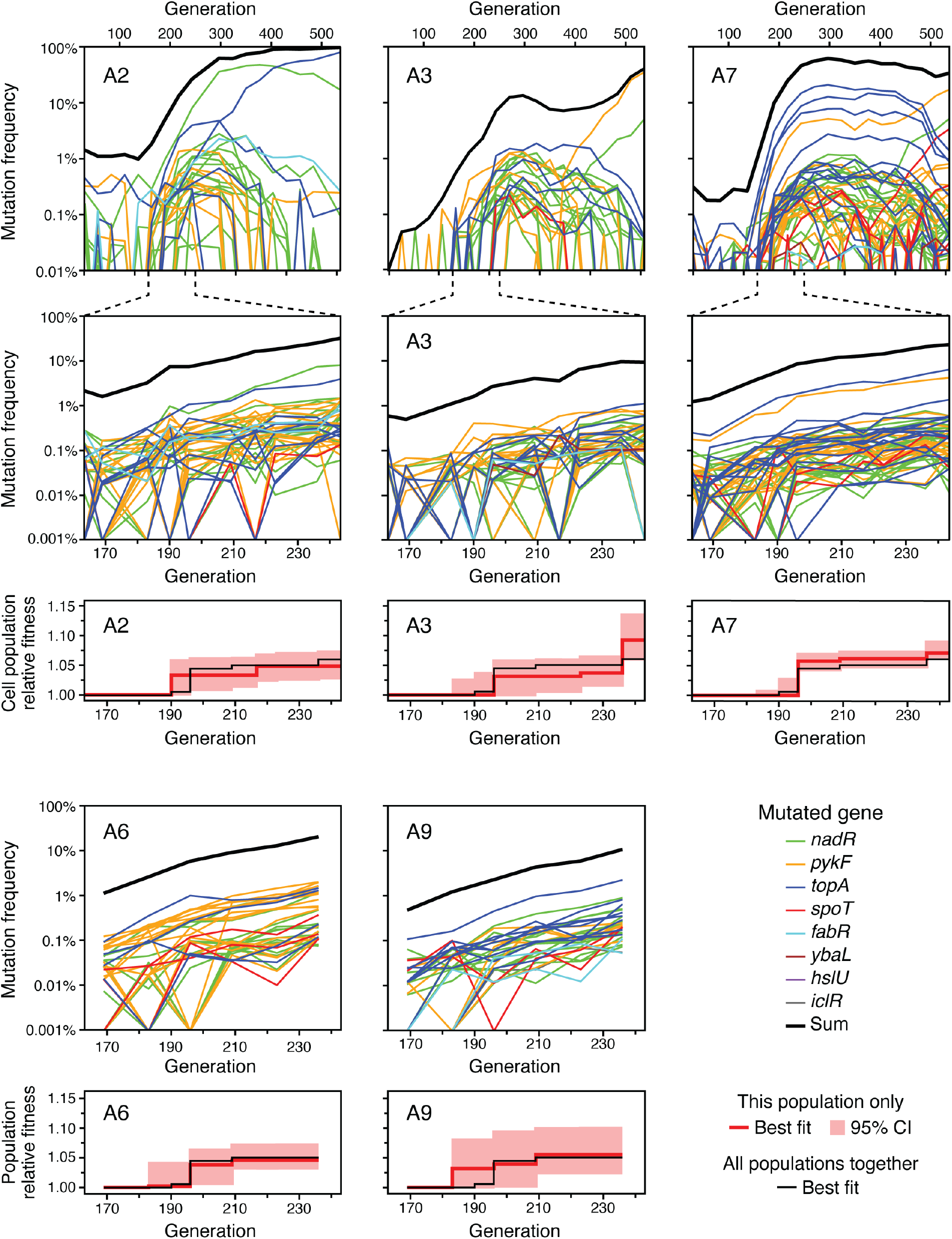
Frequency trajectories of mutations in the remaining populations. The same plots described in Figure 2C-E for population A1 are shown for populations A2, A3, and A7 (top three sets of panels). For populations A6 and A9, sequencing was only performed at time points during the selective sweep window so only the plots corresponding to Figure 2D-E are shown (bottom two sets of panels).

Mutation trajectories in all four populations exhibited a burst of genetic diversity in the targeted genes followed by loss of this diversity. The initial dynamics are expected to be largely driven by new genotypes that each evolve a single beneficial mutation very early in the experiment. If their descendants escape stochastic loss, they will gradually increase in frequency over the first few hundred generations as they outcompete the ancestral genotype. Once the population becomes dominated by these first-step mutants, their frequency trajectories plateau because of clonal interference: they are now mainly competing against one another and are relatively evenly matched. In populations A1, A2, and A7, the total frequencies of the mutations we identified sums to 50-62% at generation 270, indicating that each population is mostly composed of genotypes with a single mutation in one of the focal genes. We recovered less of the initial beneficial mutation diversity in population A3 where this sum was only 13%.

After around 300 generations, there is a steady decline in the frequencies of most mutations in the eight targeted genes. At this point, new more-fit genotypes that have evolved begin to exert their influence and displace the genotypes that we initially tracked. Many of the most successful new genotypes are descended from cells that already had a mutation in one of the targeted genes. In these cases, the original mutations serve as markers for the further expansion of these subpopulations after a period during which their frequencies stagnate or decrease, but the new beneficial mutations responsible for this further increase in fitness are outside of the genomic regions we surveilled. The converse situation, in which a beneficial mutation in one of the eight focal genes appears in a cell with an untracked beneficial mutation elsewhere in the genome, also occurs in a few cases. Most strikingly, a mutation in *pykF* that only appears after 300 generations in population A3 rapidly increases in frequency and becomes dominant, strongly suggesting that it appeared in a genetic background with a prior, unknown beneficial mutation.

### Selection coefficients can be inferred from initial mutation trajectories

We next sought to calculate the fitness benefits of individual mutations by tracking how rapidly their frequencies rose early in the experiment when they were largely competing versus the ancestral genotype because all new mutations in the population were still rare. To that end, we sequenced six populations at ~13-generation increments from 133 to 213 generations (**Fig. 2D**, **Fig. 3**). We were able to track a total of 240 mutations as they gradually increased in frequency during this critical time window. In the four populations we had already sequenced (A1, A2, A3, and A7), these mutations included 120 of the 181 previously found in the complete time course data spanning 500 generations and 54 additional mutations that were not detected in the complete time course data. We also identified 66 new mutations in the window time courses of the two populations that had not been previously sequenced (A6 and A9). Of these 240 mutations, 93.3% occurred in just three of the eight targeted genes: *nadR, pykF,* and *topA* **(Fig. 4A)**.

**Figure 4.**
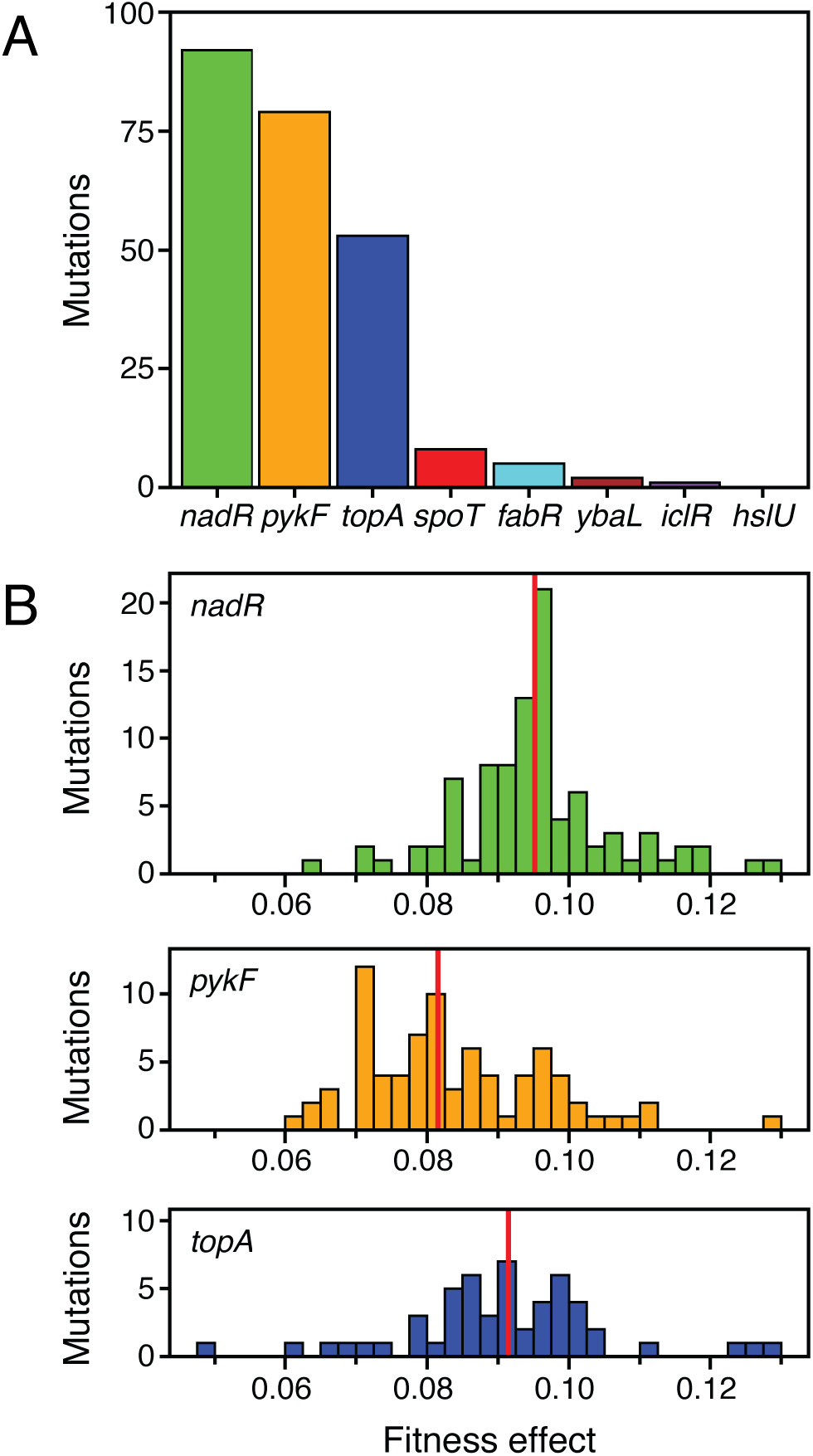
Characteristics of beneficial mutations in the initial selective sweep. (A) Total number of beneficial mutations in each targeted gene identified in the window time courses from 133 and 213 generations for all six profiled *E. coli* populations. (B) Distribution of selection coefficients of beneficial mutations determined from the window time courses in the three genes that were the dominant targets of selection. Vertical red lines show the mean of each distribution.

We were able to estimate the fitness effect of each of the beneficial mutations predicted in the window time courses by fitting a binomial logistic model to the counts of reads supporting the variant versus reference sequences over time. In all populations, there is initially a log-linear increase in the frequency of each mutation as the first wave of evolved cells, nearly all of which are expected to have just one of these beneficial mutations, competes against a population that is still almost entirely cells with the ancestral genotype. Then, there is a deceleration in the rate at which the frequencies of the new mutations increase around generation 196 that coincides with the onset of clonal interference. Genotypes with beneficial mutations begin to make up a sizable proportion of the population at this point, and making it necessary to account for how they are increasingly competing against one another to estimate the fitness effect.

We accounted for clonal interference by adding a stepwise increase in the average fitness of the entire cell population over time as an additional set of parameters to the binomial logistic model (**Fig. 2E**, **Fig. 3**). That is, we estimated how the population fitness as a whole was changing from the deceleration in the trajectories of the subset of mutations that we tracked in the targeted genes. Because overall population fitness dynamics are highly reproducible from population to population in the LTEE conditions (Lenski et al., 1991), we used one consensus population fitness increase fit from all tracked mutations in all six sequenced populations to correct our estimates of individual mutation fitness effects for clonal interference. Most of the increase in population fitness occurs rapidly in a single step during the interval spanning 196-209 generations. This rapid change followed by stasis may seem at odds with the continuing increase in the trajectories of many beneficial mutations. However, this type of stepwise increase is a typical result of clonal dynamics in models and experiments (Gerrish and Lenski, 1998; Lenski et al., 1991). It could reflect the influence of many mutations with small fitness effects, no one of which reaches an observable frequency, peaking and then being outcompeted by the most beneficial mutations that we are able to track, for example.

The mean fitness effect that we inferred for the 240 tracked mutations in all six populations was 9.00% with a standard deviation of 1.33%. Although the distributions of the fitness effects estimated for mutations in *nadR*, *pykF*, and *topA* overlap **(Fig. 4B)**, there was a significant stratification among these genes. Mutations in *nadR* were 0.44% more beneficial than mutations in *topA,* on average, and this difference was significant (*p* = 0.022, one-tailed Mann–Whitney U test). In turn, mutations in *topA* were 0.70% more beneficial than those in *pykF* (*p* = 0.00046, one-tailed Mann–Whitney U test). The fitness effects of the 16 mutations in the other genes (*spoT, fabR, ybaL,* and *iclR*) were not significantly different from the those of the 224 mutations in *nadR, pykF,* and *topA* (*p* = 0.33, two-tailed Mann–Whitney U test). Thus, highly beneficial mutations are possible in these genes as well, but they apparently occur at a much lower rate relative to that at which similarly beneficial mutations are generated in *nadR*, *pykF*, and *topA*.

### Beneficial mutations reveal different signatures of selection on gene function

Of the 301 mutations that we were able to track in complete or window time courses, 272 were in the *nadR, pykF,* or *topA* genes. These large sets of highly beneficial mutations in these genes gave us the statistical power to test for several signatures of molecular evolution to predict what types of changes in the function of each gene improved *E. coli* fitness in this environment. Each of the three genes exhibited a distinct spectrum of beneficial mutations (**Fig. 5**). In some cases, different types of mutations were also unevenly distributed throughout the sequences of these three commonly hit genes and had noticeably different effects on bacterial fitness (**Fig. 6A**).

**Figure 5.**
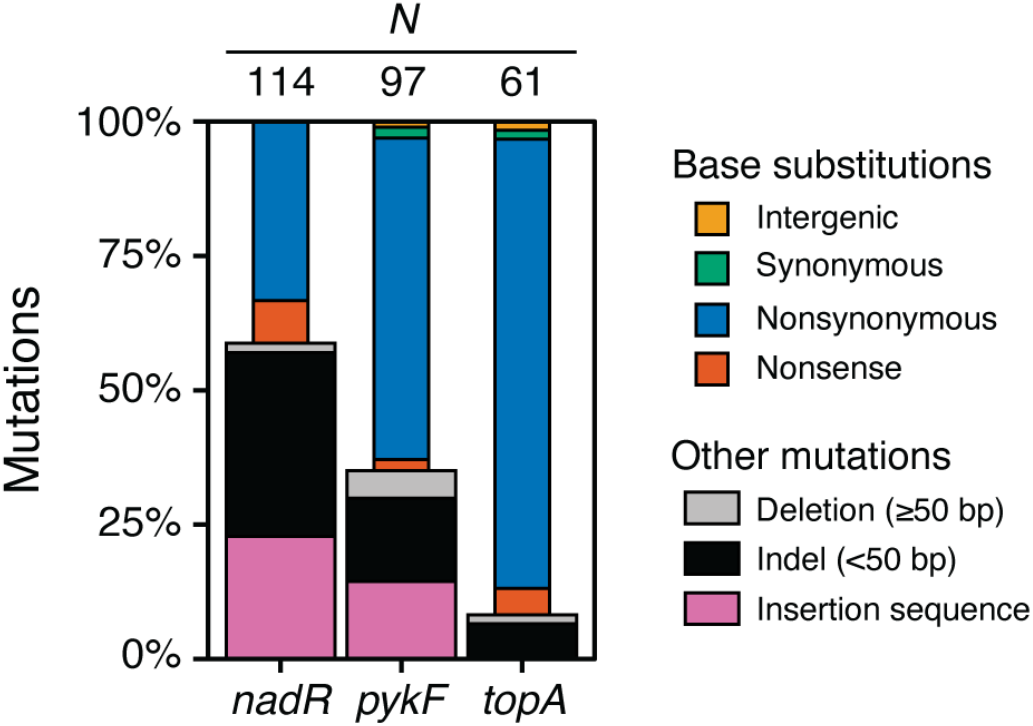
Spectra of early beneficial mutations in *nadR, pykF,* and *topA.* These three genes were the dominant targets of selection during the evolution experiment among the eight genes profiled for beneficial mutations. The bars include mutations identified in each of the six sequenced populations in its window time course, complete time course, or both. The total number of mutations identified in each gene is indicated above its column.

**Figure 6.**
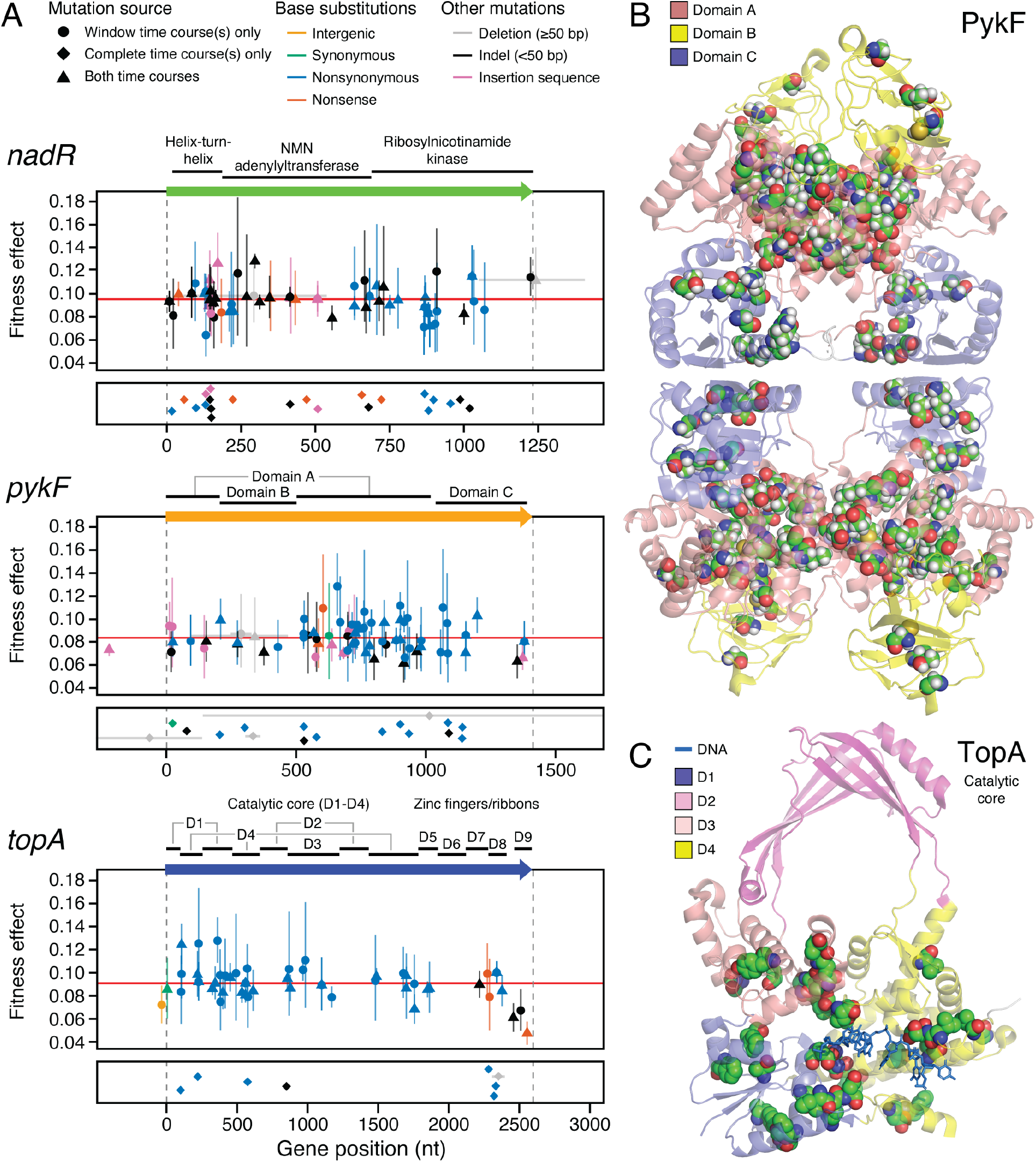
Mutations in the three genes that were the dominant targets of selection. (A) Nucleotide positions and properties of all mutations found in each of the three genes that were the dominant targets of selection during the evolution experiment. Colors represent the type of mutation. Symbols indicate whether each mutation was detected in the window time course, the complete time course, or both. The upper panel for each gene, shows the fitness effects estimated for mutations in that gene from the window time courses. Error bars are 95% confidence limits. When the same genetic change was detected in the window time courses for multiple populations the trajectories were combined to estimate one overall fitness value and confidence limit for all of those mutations (see **Methods**). Thus, the symbols and error bars for these mutations overlap in these cases. The reading frame of the gene is shown above this panel with protein domains labeled. Vertical dashed grey lines represent the start and end of each gene. Horizontal grey lines show the extent of large deletions within the pictured region. Horizontal red lines represent the average fitness effects for all mutations in a gene. The lower panel for each gene shows mutations that were only detected in the complete time courses and therefore do not have an estimated fitness effect. Symbols in this panel are randomly jittered in the vertical direction to improve their visibility. (B) Structural context of mutations in PykF. Sites of nonsymonymous mutations are highlighted by showing space-filling models of the substituted amino acid residues. All four subunits of the PykF homotetramer are shown. (C) Structural context of mutations in the catalytic core of TopA. Sites of nonsymonymous mutations are highlighted by showing space-filling models of the substituted amino acid residues. Only domains D1-D4 are present in the structure. The DNA strand interacting with TopA is shown as a stick model.

The *E. coli nadR* gene has three distinct functions related to NAD biosynthesis: (1) the N-terminal domain is a helix-turn-helix that binds to DNA so that it can act as a negative transcriptional regulator of NAD salvage and transport pathways; (2) the internal domain is an NMN adenylyltransferase (Raffaelli et al., 1999); and (3) the C-terminal domain is predicted to have ribosylnicotinamide kinase activity (Kurnasov et al., 2003). Large deletions, frameshifts from small insertions or deletions (indels), insertions of transposable insertion sequence (IS) elements, and base substitutions creating stop codons dominate the *nadR* mutational spectrum (**Fig. 5**). These disruptive mutations, which are expected to result in complete loss of gene function, are significantly overrepresented versus nonsynonymous base substitutions in the first two domains of the gene compared to the remainder (13.7 odds ratio, *p* = 1.2 × 10^-8^, one-tailed Fisher’s exact test) (**Fig. 6A**). Yet, there was not a significantly greater selection coefficient for disruptive mutations compared to nonsynonymous mutations overall (*p* = 0.063, one-tailed Mann–Whitney U test). These results suggest that complete inactivation of *nadR* yields the maximum benefit possible for a mutation in this gene. Disrupting all three of its distinct functions does not appear to be necessary for achieving this full benefit. Consistent with this prediction, deletion of *nadR* has been shown to be highly beneficial in the very similar environment of the LTEE (Barrick et al., 2009).

Pyruvate kinase 1 (*pykF*) catalyzes the final step of glycolysis, transferring a phosphate group from phophoenolpyruvate (PEP) to ADP to generate pyruvate and ATP. It is a key enzyme in regulating glycolytic flux (Kochanowski et al., 2013; Siddiquee et al., 2004). We observed an intermediate representation of disruptive mutations in *pykF,* fewer than in *nadR* but more than in *topA* (**Fig. 5**). Interestingly, nonsynonymous base substitutions in *pykF* tend to have a larger selection coefficient than disruptive mutations (*p* = 0.00031, one-tailed Mann–Whitney U test) (**Fig. 6A**). This finding is in agreement with a study of various *pykF* alleles that arose in the LTEE which found that nearly all *pykF* point mutations were more beneficial than deletion of the *pykF* gene, both in the ancestor and in evolved genetic backgrounds (Peng et al., 2018). PykF forms a homotetramer in which each polypeptide folds into three structural domains (Donovan et al., 2016; Mattevi et al., 1995). The central domain A forms the active site at the interface with domain B and the binding site for the allosteric effector fructose 1,6-bisphosphate at the interface with domain C. The nonsynonymous mutations that we observed are more concentrated than expected in domain A versus the other structural domains based on their relative lengths in the gene sequence (*p* = 0.0018 one-tailed binomial test) (**Fig. 6B**). Overall, these results suggest that complete inactivation of *pykF* is highly beneficial in the environment of our evolution experiment, but mutations that alter its activity—likely in ways that reduce glycolytic flux—are even more so. It has been suggested that reducing *pykF* activity is beneficial in the similar glucose-limited conditions of the LTEE because this allows more PEP to be diverted to power import of glucose into cells via the phosphotransfer system (Woods et al., 2006).

DNA topoisomerase I (*topA*) relaxes negative supercoiling introduced into the chromosome by replication and transcription (Massé and Drolet, 1999). The mutations we observed in *topA* are almost exclusively single-base substitutions (**Fig. 5**), suggesting that modulating the activity of this enzyme provides a fitness benefit. Indeed, complete loss of *topA* function is lethal to *E. coli* without compensatory mutations in DNA gyrase (Dinardo et al., 1982; Pruss et al., 1982). The structure of *E. coli* TopA consists of four N-terminal domains (D1-D4) that make up the catalytic core and five C-terminal zinc finger and ribbon domains (D5-D9) (Tan et al., 2015). The few out-of-frame indels and the large deletion that we observe truncate TopA within domains D7-D9, which interact with single-stranded DNA and RNA polymerase but are not critical for catalysis. Considering only the catalytic core, we find that nonsynonymous mutations are more concentrated in domains D1 and D4 versus D2 and D3 than expected from their relative sizes (*p* = 0.00068, one-tailed binomial test) (**Fig. 6C**). D1 and D4 together form the ssDNA binding groove leading to the active site, and D1 also forms part of the active site at its interface with D3 (Perry and Mondragón, 2003). Several base substitutions in *topA* have been shown to increase positive supercoiling in evolved LTEE strains (Crozat et al., 2005, 2010). The exact reason that this change in supercoiling is beneficial is unknown, but it may be linked to increasing the expression of ribosomal RNAs (Crozat et al., 2005), altering gene regulation responses to starvation or other stresses (Crozat et al., 2010), and/or increasing the expression of genes in divergently transcribed operons (Houdaigui et al., 2019).

### Recurrent beneficial mutations do not have greater fitness effects

We observed many examples of exact genetic parallelism. That is, the same mutation occurred and reached high frequency in different experimental populations. Each of these *E. coli* populations was founded from single cells, so we can conclude that these recurrent genetic changes are due to independent mutational events. We observed a total of 252 distinct genetic changes across all eight profiled genes and 31 of these were found in more than one population.

While no single genetic change was detected in all six populations, 2, 2, 8 and 19 changes were detected in 5, 4, 3, and 2 populations, respectively. Most of these were in the three genes that were the main targets of selection (*nadR, pykF,* and *topA*), but one that occurred in three populations was in *fabR.* These mutations may be recurrent because they have a higher fitness benefit than other mutations, occur at a higher rate than other mutations, are more easily detected in the sequencing data, or due to some combination of these factors. We had a fitness estimate for each of the 31 recurrent mutations from tracking cells with that genetic change in at least one of the window time courses and had 167 fitness measurements for mutations that were observed in only one population. The recurrent mutations had a 0.12% greater fitness effect, on average, compared to the singleton mutations, but this difference was not significant (*p* = 0.25, one-tailed Mann–Whitney U test). Thus, it is unlikely that many cases of exact genetic parallelism are due to these mutations being more beneficial than others in our dataset.

## Discussion

We examined bacterial evolution during the initial stages of clonal competition when there is a burst of beneficial genetic diversity as many new subpopulations with different mutations evolve and begin to displace the ancestral genotype. We focused on eight genes known to accumulate adaptive mutations in the >70,000 generation Lenski long-term evolution experiment (LTEE) with *E. coli* that used nearly the same environment as our experiments. The only difference was that we added four times as much of the limiting nutrient (glucose). By combining Illumina sequencing using unique molecular identifiers for error correction, hybridization-based capture of DNA encoding these genes, and dense temporal sampling, we were able to identify more beneficial mutations and track them at much lower frequencies than is possible with standard metagenomic sequencing. We detected a total of 301 mutations in the focal genes: 181 in the complete time courses of four populations and 240 in the window time courses of these populations and two others, with 120 mutations overlapping between the two sets.

By densely sampling and deeply sequencing *E. coli* populations, we were able to characterize many beneficial mutations that never reach the detection limits of standard Illumina sequencing before they become casualties of clonal interference. Only 13 of the 181 mutations we detected in the complete time courses ever achieved a frequency of 5% or more, which can be reliably distinguished from noise by standard metagenomic sequencing, and only seven were this common for 100 or more generations, such that they were likely to be detected by a typical time-sampling scheme. Considering both the complete and window time courses we characterized 241 and 42 mutations that never reached 1% or 0.1% thresholds, respectively, at any of our sampled time points. Our success in recovering rare variants meant that we discovered more examples of beneficial mutations in the three commonly mutated genes (*topA*, *pykF*, and *nadR*) than have been reported in many prior studies of the evolution of the twelve LTEE populations through 60,000 generations of evolution (Barrick et al., 2009; Deatherage et al., 2015; Good et al., 2017; Ostrowski et al., 2008; Tenaillon et al., 2016; Woods et al., 2006). These large sets of mutations enabled us to identify distinct molecular signatures of adaptation in each protein.

We profiled evolution driven by mutations in eight genes known to be targets of selection in the LTEE. Mutations in four of these (*topA*, *pykF, spoT,* and *fabR*) reach high frequencies within the first 1,000 generations of the LTEE in multiple populations (Deatherage et al., 2015; Good et al., 2017). Mutations in the other four (*hslU, nadR, ybaL,* and *iclR*) are also common in the LTEE, but they typically occur later (often within the first 2,000 to 10,000 generations) (Good et al., 2017; Tenaillon et al., 2016). Nearly all mutations in these genes in our evolution experiment were in *topA*, *pykF*, and *nadR*, but we also found multiple mutations that were similarly beneficial in *spoT, fabR,* and *ybaL.* Mutations in *nadR* were more widespread than expected in our experiment and may be more likely to completely disrupt its function than beneficial alleles that evolve in the LTEE (Ostrowski et al., 2008). Mutations in *spoT* and *fabR* were rarer than expected from the LTEE. One possible explanation for these slight differences is the increased concentration of glucose in our experiment compared to the LTEE. These minor deviations are also reminiscent of how changing a different aspect of the environment (temperature) re-focuses the mutations of largest benefit that succeed early onto different subsets of genes, nearly all of which eventually accumulate beneficial mutations later in the LTEE environment, in related evolution experiments (Deatherage et al., 2017; Tenaillon et al., 2012). Alternatively, we cannot rule out that mutations with similar fitness effects are still possible in the rarely mutated genes but that there are so many fewer possibilities for these mutations that they were not often sampled in our evolving populations. Despite these subtle differences between the LTEE and our experiment, we were still able to account for majority of the genetic variation present in three of the four populations that we profiled over the entire 500 generations by analyzing evolution in the eight candidate genes.

We also wanted to understand to what extent we gained early warning of driver mutations by deeply profiling evolution in genes we expected to be under strong selection. In general, we were able to begin tracking most mutations when they were above a frequency of 0.01%. This level of profiling enabled us to first detect mutations an average of 69, 150, and 290 generations before they surpassed frequencies of 0.1%, 1%, and 5%, respectively. Under the conditions of our experiment these intervals take roughly 10, 23, and 44 days, respectively; so, even though we made these predictions retrospectively, there would have been sufficient time to complete the DNA isolation, library preparation, sequencing, and analysis steps quickly enough for this approach to give early warning of specific genetic variants driving evolution of these populations. The amount of lead time becomes disproportionately longer at higher frequencies due to clonal interference between beneficial mutations. The chances and timescales of earlier detection are expected to increase even more when there are ecological interactions or spatial structure that further slow the takeover of new variants, as has been demonstrated and discussed in other microbial evolution experiments (Baym et al., 2016; Frenkel et al., 2015; Traverse et al., 2013).

A further prediction is that the genes in which we observe early, but unsuccessful beneficial mutations will sustain mutations again and again until they are successful in a population’s evolutionary future. This prediction is limited by the nature of epistatic interactions. In the LTEE and other microbial evolution experiments, diminishing returns epistasis dominates between beneficial mutations in different genes (Chou et al., 2011; Khan et al., 2011; Kryazhimskiy et al., 2014; Wei and Zhang, 2019; Wiser et al., 2013). That is, mutations in one gene that improve the fitness of the ancestor tend to still be beneficial to evolved genotypes containing beneficial mutations in other genes, just less so than when those other mutations are not present. Subpopulations with mutations in both *nadR* and *pykF* evolve by 20,000 generations in all 12 LTEE populations, and cells that also contain a mutation in *topA* are found in six of the LTEE populations at this point (Tenaillon et al., 2016). By this time, mutations in *ybaL* and *spoT* are also found in nine and six LTEE populations, respectively. So, for five of the six genes in which we detected multiple mutations in the initial burst phase, it is likely that nearly all of them would have eventually accumulated beneficial mutations if we continued our experiment.

The other three genes (*fabR*, *iclR,* and *hslU*) likely represent other scenarios. Mutations in *fabR* transiently appear within the first 2,000 generations of the LTEE (Deatherage et al., 2015). They interact unfavorably with beneficial mutations in *spoT* and other genes, such that a *fabR* mutation essentially precludes further adaptation by mutating the other set of genes and vice-versa (Deatherage et al., 2015; Woods et al., 2011). We detected 9 mutations in *fabR,* which was more than the five we observed in *ybaL*. However, we predict that *fabR* mutations are unlikely to re-emerge and be successful in the future of these populations because of their negative interactions with other beneficial mutations. On the other hand, we detected only a single mutation in *iclR* and a single mutation in *hslU.* Of the 12 LTEE populations, 11 have sizable subpopulations with mutations in *iclR* and 11 have mutations in *hslU* by 20,000 generations, which makes them more common than mutations in *spoT* and *ybaL* in the long run. Therefore, mutations in *iclR* and *hslU* appear to either require the presence of mutations in other genes to become highly beneficial or may not be able to experience any mutations that are beneficial enough to make them competitive early on in our experiment.

The nature of epistasis and the limits that it imposes on predicting the future evolution of a cell population could be further probed using our approach in several ways. One could repeat the evolution experiment beginning with genotypes containing different first-step beneficial mutations as starting points to more finely map the fitness landscape. One could also interrogate the diverse collections of cells containing different beneficial alleles that we have evolved, by taking the 150-generation populations and further evolving them under different conditions to map genotype by environment effects, for example. Such experiments might also reveal latent beneficial mutation in other genes (e.g., *iclR* and *hslU*) that were able to outcompete the ancestor in our experiment but remained below the detection limit because they were not as beneficial as mutations in *topA*, *pykF*, and *nadR* in this environment. There is precedent for changes in the environment deflecting selection to different subsets of the same genes. In an offshoot of the LTEE that began with a clone that had *spoT, topA,* and *pykF* mutations, selection was focused on either *hslU, iclR,* or *nadR* depending on changes in temperature (Deatherage et al., 2017).

Alternative and complementary methods exist for deeply profiling the evolutionary possibilities inherent in the fitness landscape of a cell, i.e., its evolvome. We tracked spontaneous beneficial mutations within targeted genome regions, or a portion of what one could more specifically describe as the adaptome (Ryall et al., 2012). Amplicon sequencing can also capture mutations in a subset of the genome with deep coverage (Chubiz et al., 2012; Fischer et al., 2017; Hong et al., 2018). We used hybridization-based enrichment, which did not require any experimental optimization for different targets and is less likely to introduce biases in inferring the frequencies of mutations when IS insertions and large indels that change amplicon sizes are present. Similar procedures have been used to detect mutations associated with cancer when they are only present at very low frequencies in circulating tumor DNA found in patient blood samples (Newman et al., 2014, 2016). Tracking the frequencies of barcoded cells and their progeny has been used to characterize the statistical properties of much larger collections of naturally occurring beneficial mutations and when they are much rarer within cell populations (Levy et al., 2015; Venkataram et al., 2016). However, one must barcode individuals in the population to apply this method, which may be difficult in certain cell types or in clinical samples, and additional genome sequencing after an experiment is completed is required to discover the identities of the beneficial mutations linked to barcodes. Other methods such as deep-mutational scanning (Fowler and Fields, 2014) or CRISPR-enabled trackable genome engineering (Garst et al., 2017) can simultaneously interrogate large libraries of mutants to map evolvomes. However, since they artificially construct variant libraries, they do not necessarily provide information about which genetic variants are accessible by spontaneous mutations and would therefore be expected to contribute the most to a cell’s adaptome.

Exhaustively mapping paths that clonal evolution is likely to follow is of particular interest and utility in systems that evolve repeatedly from a defined starting point. These range from bioreactors that are seeded with the same strain in different production runs to lung infections in cystic fibrosis patients that start from similar, but not identical, opportunistic pathogens. The ability to identify mutations in key genes while they are still very rare may also be used to improve the early detection of emerging drug resistance in other human infections and cancer. The evolutionary dynamics will be more complex in many of these systems, but they may also unfold more slowly. For example, biofilm formation and the necessity of invading already colonized niches will slow the dynamics of competition. This potentially makes the therapeutic window for detecting incipient evolution by profiling the adaptome even greater.

## Materials and Methods

### Evolution experiment

Strains and growth conditions are derived from the Lenski long-term evolution experiment (Lenski and Travisano, 1994; Lenski et al., 1991). Nine colonies of *E. coli* B strain REL606 and nine of strain REL607 were selected at random. Each was used to inoculate a separate flask containing 10 mL of Davis Minimal (DM) media supplemented with 100 μg/L glucose (DM100). This is a slightly higher concentration of glucose than the 25 μg/L glucose (DM25) used in the LTEE, but still well below the ~1000 μg/L concentration at which nutrients other than glucose begin to limit growth in this medium. We used this higher glucose concentration to ensure we had a sufficient number of cells for sequencing and archiving. These initial cultures were grown overnight at 37°C with orbital shaking over a one-inch diameter at 120 RPM. Approximately 30 generations of growth occurred starting from the initial single cell that gave rise to each colony until saturation of these cultures. Next, 10 mL of fresh DM100 were inoculated with 50 μL of one REL606 culture and 50 μL of one REL607 culture for overnight growth in the same conditions. The remaining culture was archived at −80°C with 2 mL dimethyl sulfoxide (DMSO) added as cryoprotectant. Daily transfer of 100 μL of overnight culture to 10 mL of fresh DM100 and archival of the remaining culture volume in the same way continued through 75 daily transfers. Periodically 1 μL of culture was diluted 10,000-fold in sterile saline and plated on tetrazolium arabinose (TA) agar to allow growth of ~200 colonies. REL606 and REL607 differ by a mutation in an arabinose utilization gene that makes REL606 (Ara^-^) colonies red and REL607 (Ara^+^) colonies pink (Lenski et al., 1991). The ratio of red to pink colonies was used to monitor these populations for selective sweeps (Hegreness and Kishony, 2007; Woods et al., 2011).

### DNA isolation and library preparation

Genomic DNA (gDNA) was isolated from frozen population samples by first thawing each 15 mL conical tube on ice. Of the ~12 mL total volume of culture plus cryoprotectant, 1.2 mL was transferred to a 2 mL cryovial and refrozen. The remaining ~10.8 mL was centrifuged at 6,500 × g at 4°C for 15 minutes. The resulting cell pellets were transferred with a volume of remaining solution to 1.7 mL Eppendorf tubes. Then, gDNA was isolated using the PureLink Genomic DNA Mini kit (Life Technologies). For each sample, 1 μg of gDNA was randomly fragmented on a Covaris S2 focused-ultrasonicator.

Illumina libraries were constructed using the Kappa Biosystems LTP Library Preparation Kit with the following modifications. End-repaired, fragmented DNA was T-tailed (rather than A-tailed) in a 50 μl reaction including 10 mM dTTP and 5 units of Klenow fragment, exo^-^ (New England Biolabs). Illumina adapters containing 12-base unique molecular identifiers were ligated to the T-tailed fragments as previously described (Schmitt et al., 2012), except full-length adapter sequences containing unique external sample barcodes were directly ligated to the T-tailed dsDNA inserts to reduce the risk of cross-contamination between samples. The full list of DNA sequence adaptors used is provided in **Table S1**.

### Probe design and target capture

Oligonucleotide probes consisting of 60-base xGen Lockdown probes (Integrated DNA Technologies) were designed to tile across each of the eight genes of interest including upstream promoter elements. Probes for each gene were compared to the entire *E. coli* B strain REL606 reference genome (GenBank: NC_012967.1) (Jeong et al., 2009) using BLASTN (Camacho et al., 2009). The starting positions of all probes in a set were shifted by one base at a time until every probe had only a single significant predicted binding location (match with E-value < 2×10^-5^). The sequences of the final set of 242 probes are provided in **Table S2**.

Capture was performed using a SeqCap EZ Exome Enrichment kit v3.0 (NimbleGen) with several modifications to the protocol. First, 18 to 20 population samples with different sample barcodes were pooled together in a single capture reaction that contained a total of 1 μg of library DNA from all samples, 1 mmol of a universal blocking oligo, and 1 mmol of a degenerate sample barcode blocking oligo. The sequences of these blocking oligos are provided in **Table S3**. Second, after hybridization for 72 h, DNA fragments hybridized to the biotinylated probes were recovered using MyOne Streptavidin C1 Dynabeads (Life Technologies). Third, a final 8-cycle PCR step was performed with HiFi Hotstart DNA Polymerase (Kappa Biosystems).

### Sequencing and read processing

Paired-end 101- or 125-base sequencing of the final libraries was performed on an Illumina HiSeq 2000 at the University of Texas at Austin the Genome Sequencing and Analysis Facility (GSAF). Read sequences have been deposited into the NCBI Sequence Read Archive (PRJNA601748). Raw reads were used to generate Consensus Sequence Reads (CSR) using custom Python scripts that carried out the following steps. First, the beginning of each read was evaluated for the presence of the expected 5’-end tag, consisting of a random 12-base unique molecular identifier (UMI) followed by four fixed bases (5’-N_12_CAGT). Read pairs lacking the correct 5’-end tag on either read were discarded. Across all samples, 80.2% of read pairs had both UMIs. For remaining read pairs, the UMIs found on each end were concatenated to create a 24-base dual-UMI that uniquely identifies the original DNA fragment that was amplified and sequenced. To group all reads corresponding to the same initial DNA molecule, a FASTA file of all dual-UMIs was used as input into the *ustacks* program from the Stacks software pipeline (Version 1.48) (Catchen et al., 2013) with the following options: a single read was sufficient to seed a stack, a single mismatch within the dual-UMI was allowed in assigning a read to a stack, secondary reads and haplotypes were disabled, and stacks with high coverage were preserved. Then, CSRs were generated for all dual-UMI groups sequenced at least twice by taking the straight consensus of all reads that were merged into that stack. If no base exceeded 50% frequency at a given position in this set of reads, then that base was set as unknown (N). Of the read pairs with valid dual-UMIs, 41.6% were incorporated into consensus reads across all samples. The average number of dual-UMI read pairs utilized to create each consensus read was 2.46, which gives an overall yield of consensus reads per read in a pair with a valid dual-UMI 16.9%.

### Variant calling

We used the *breseq* pipeline (Barrick et al., 2014; Deatherage and Barrick, 2014; Deatherage et al., 2015) (version 0.26.0) to call single-nucleotide variants (SNVs) and structural variants (SVs) from the CSRs. We divided the genome sequence of the ancestral *E. coli* REL606 strain into two types of reference regions for mapping in this analysis. The eight regions of the genome tiled with probes—extended with hundreds of bases of flanking sequence on both sides—were input as “targeted” sequences, and the remainder of the genome with the identical eight regions masked to degenerate N bases was supplied as a “junction-only” reference (to which reads are mapped without variant calling). All 116 samples were analyzed using *breseq* in polymorphism prediction mode with all bias, minimum allele frequency, and read-count filters disabled. Evidence items in the Genome Diff (GD) files for all samples were combined using the *gdtools* utility program to generate a single merged GD file with each piece of evidence listed a single time, regardless of how many times it was detected in different samples. We then re-ran *breseq* using the same parameters except that this GD file was supplied as an input user-evidence file to force output of variant and reference information for these putative variants in every sample. Then, we extracted the number of variant reads supporting each putative variant allele and the total number of reads at that reference location from the GD file output by *breseq.* Subsequent statistical tests and fitting steps were performed in R (version 4.0.0) (R Core Team, 2016) using the ggplot2 package for data visualization (Wickham, 2016). Scripts and data files for this analysis are available in GitHub (https://github.com/barricklab/adaptome-capture).

Since this original analysis was conducted at the level of *breseq* evidence (i.e., single columns of read pileups on the reference genome or instances of new sequence junctions), we next merged sets of observations that were consistent with a single mutational event when they also had frequency trajectories that tracked together. To identify these candidates for merging, we analyzed each of the six window (generation 133 to 213) and four complete (generation 0 to 500) time courses separately. We only considered mutations that exceeded a threshold frequency of 0.03% at some time during each time course as candidates for merging.

Read alignment (RA) evidence items were merged when they were located within 6 base pairs of one another and within a normalized Canberra distance of 0.1 between the vectors of their frequency observations across all of the time points in a dataset. All RA evidence pairs of this kind were found to co-occur in the same sequencing reads. For these cases, the read counts for the first linked mutation were used to represent the entire event. For example, if a deletion of three base pairs was predicted by missing bases at positions x, y, and z; then the frequency of missing the first base (x) was assigned to the entire three-base deletion mutation.

For new junction (JC) evidence we performed the same merging procedure but allowed linked mutations to be within a larger window of 20 base pairs and within a normalized Canberra distance of 0.5. JC pairs passing these criteria were only merged if they were also consistent with an IS-element insertion in terms of their relative orientation and spacing. In this case the variant and total read counts were added together for the two different junctions, as the junctions on each side of the inserted IS element provide independent information for estimating the frequency of this type of mutation. We allowed unpaired JC evidence passing the filters to also predict IS element mutations. This situation may indicate that there was an IS-mediated deletion between an element that inserted within the gene and another element from the same family located outside of the targeted region or more complex chromosomal rearrangements involving a newly inserted IS element (Raeside et al., 2014).

### Time course filtering and fitness effect estimation

After merging evidence of genetic variants into lists of putative mutations, we further eliminated some of these from consideration using several filtering steps. For the complete time courses, we first required that non-zero frequencies be observed for a mutation in samples from at least two different time points. We next applied a filter to eliminate spurious variants that can be recognized as arising from systematic sequencing or alignment errors because they do not exhibit the correlated changes in frequency over time expected for the frequency trajectories of real mutations (Lang et al., 2013). Specifically, we required that the time-series of estimated frequencies for a mutation over all analyzed time points have an autocorrelation value ≥ 0.55.

For the window time courses, we eliminated putative mutations for which there was great uncertainty in the estimated fitness effect or evidence that its trajectory reflected multiple beneficial mutations occurring in the same genome. Specifically, we required that a mutation was first observed at generation 196 or earlier and that its estimated frequency was ≥ 10 ^4^ in every sample that was sequenced from generation 223 to 243. Then, we fit a binomial logistic model with slope and *y*-intercept terms to the time courses of counts of variant and reference (total minus variant) observations for each mutation. We used a negative offset in the model of the number of generations up to each time point so that the slope represents one plus the selection coefficient that is characteristic of the subpopulation with that mutation. We filtered out any mutations for which this fit had an AIC < 200, a *p*-value for the slope differing from zero of > 0.005, or a *y*-intercept < −20. The fitness effects that we report for mutations are the selection coefficients fit from the model divided by the natural logarithm of two so that they are expressed per generations of binary cell division. One plus the fitness effect is the relative fitness of a cell with that mutation. These values can be directly compared to experimental measurements of relative fitness and mutation fitness effects made using co-culture competition assays (Lenski et al., 1991; Tenaillon et al., 2016).

This procedure for determining fitness effects assumes that the trajectories reflect competition purely against the ancestral strain. However, we detected a consistent deviation from linearity for all mutation trajectories in the window time courses after generation 196. The rates at which the frequencies of all mutations were increasing decelerated, indicating that the overall population fitness had improved to a degree that it reduced their effective advantage versus their competitors. To account for this change we fit additional parameters defining a stepwise increase in the average relative fitness of the population within each interval between sequenced samples from generation 196 onward. The increase in population fitness reduces the effective time basis used in the model to determine the slope to the number of generations in each interval divided by the average relative fitness during that interval. We determined the population fitness values that minimized the AIC of this modified binomial logistic model. The figures show the best stepwise increases in population fitness between the sequenced time points from generation 196 onward fitting to the trajectories of all mutations in a given population at the same time. We performed 1000 bootstrap resamplings of the mutations in each population to estimate 95% confidence intervals on the estimated population fitness values in each interval for that population.

We combined information across multiple populations in two ways to further improve the estimates of mutation fitness effects. First, there was considerable uncertainty in the estimates of the stepwise population fitness increases for each population considered alone. Because the actual population fitness trajectories of all populations are expected to be highly similar to one another, we fit a consensus stepwise increase in population fitness over time that best improved the fits for all mutations from all populations. Second, we observed 27 cases in which the exact same change in a gene’s sequence was observed and passed our filtering criteria in the window time courses of multiple experimental populations. Because each population was started from single cells, we can be sure that these are independent observations of the same mutation. Therefore, we fit one consensus fitness effect (slope) for each of these recurrent mutations across all populations. We still allowed the *y*-intercept for each of these mutations to vary from population to population because this parameter is related to how early the mutation evolved, which is expected to be different in each replicate population.

### Protein structure analysis

Structural domains in NadR, PykF, and TopA were defined according to UniProt and papers reporting x-ray crystal structures. Mutations in PykF were mapped onto Protein Data Bank structure 4YNG (Donovan et al., 2016). Mutations in TopA were mapped onto Protein Data Bank structure 1MW8 (Perry and Mondragón, 2003). Protein structures were visualized using Pymol v2.3.5 (Schrödinger LLC).

## Acknowledgements

The authors acknowledge the Texas Advanced Computing Center (TACC) at The University of Texas at Austin for providing high-performance computing resources.

## Supporting Information

**Table S1** Adapter sequences used in DNA library preparation

**Table S2** Sequences of pulldown probes

**Table S3** Blocking oligos used to limit read-to-read binding during pulldown

## Author contributions

Conceptualization: DED JEB.

Data Curation: DED JEB.

Funding Acquisition: DED JEB.

Investigation: DED.

Methodology: DED JEB.

Software – DED JEB.

Supervision – JEB.

Visualization: DED JEB.

Writing – Original Draft Preparation: DED JEB.

Writing – Review & Editing: DED JEB.

## Data Availability Statement

DNA sequence files are available from the NCBI Sequence Read Archive (accession number PRJNA601748). Code and processed data files are available on GitHub (https://github.com/barricklab/adaptome-capture).

## References

Bainbridge, M.N., Wang, M., Burgess, D.L., Kovar, C., Rodesch, M.J., D’Ascenzo, M., Kitzman, J., Wu, Y.-Q., Newsham, I., Richmond, T. a, et al. (2010). Whole exome capture in solution with 3 Gbp of data. Genome Biol. 11, R62.

Barrick, J.E. (2020). Limits to predicting evolution: insights from a long-term experiment with *Escherichia coli*. In Evolution in Action: Past, Present and Future, W. Banzhaf, B.H.C. Cheng, K. Deb, K.E. Holekamp, R.E. Lenski, C. Ofria, R.T. Pennock, W.F. Punch, and D.J. Whittaker, eds. (Cham: Springer), pp. 63–76.

Barrick, J.E., and Lenski, R.E. (2009). Genome-wide mutational diversity in an evolving population of *Escherichia coli*. Cold Spring Harb. Symp. Quant. Biol. 74, 119–129.

Barrick, J.E., and Lenski, R.E. (2013). Genome dynamics during experimental evolution. Nat. Rev. Genet. 14, 827–839.

Barrick, J.E., Yu, D.S., Yoon, S.H., Jeong, H., Oh, T.K., Schneider, D., Lenski, R.E., and Kim, J.F. (2009). Genome evolution and adaptation in a long-term experiment with *Escherichia coli*. Nature 461, 1243–1247.

Barrick, J.E., Colburn, G., Deatherage, D.E., Traverse, C.C., Strand, M.D., Borges, J.J., Knoester, D.B., Reba, A., and Meyer, A.G. (2014). Identifying structural variation in haploid microbial genomes from short-read resequencing data using *breseq*. BMC Genomics 15, 1039.

Baym, M., Lieberman, T.D., Kelsic, E.D., Chait, R., Gross, R., Yelin, I., and Kishony, R. (2016). Spatiotemporal microbial evolution on antibiotic landscapes. Science 353, 1147–1151.

Camacho, C., Coulouris, G., Avagyan, V., Ma, N., Papadopoulos, J., Bealer, K., and Madden, T.L. (2009). BLAST+: architecture and applications. BMC Bioinformatics 10, 421.

Catchen, J., Hohenlohe, P.A., Bassham, S., Amores, A., and Cresko, W.A. (2013). Stacks: an analysis tool set for population genomics. Mol. Ecol. 22, 3124–3140.

Chou, H.-H., Chiu, H.-C., Delaney, N.F., Segrè, D., and Marx, C.J. (2011). Diminishing returns epistasis among beneficial mutations decelerates adaptation. Science 332, 1190–1192.

Chubiz, L.M., Lee, M.-C., Delaney, N.F., and Marx, C.J. (2012). FREQ-Seq: a rapid, cost-effective, sequencing-based method to determine allele frequencies directly from mixed populations. PLoS ONE 7, e47959.

Crozat, E., Philippe, N., Lenski, R.E., Geiselmann, J., and Schneider, D. (2005). Long-term experimental evolution in *Escherichia coli*. XII. DNA topology as a key target of selection. Genetics 169, 523–532.

Crozat, E., Winkworth, C., Gaffé, J., Hallin, P.F., Riley, M.A., Lenski, R.E., and Schneider, D. (2010). Parallel genetic and phenotypic evolution of DNA superhelicity in experimental populations of *Escherichia coli*. Mol. Biol. Evol. 27, 2113–2128.

Cvijović, I., Nguyen Ba, A.N., and Desai, M.M. (2018). Experimental studies of evolutionary dynamics in microbes. Trends Genet. 34, 693–703.

Deatherage, D.E., and Barrick, J.E. (2014). Identification of mutations in laboratory-evolved microbes from next-generation sequencing data using *breseq*. Methods Mol. Biol. 1151, 165–188.

Deatherage, D.E., Traverse, C.C., Wolf, L.N., and Barrick, J.E. (2015). Detecting rare structural variation in evolving microbial populations from new sequence junctions using *breseq*. Front. Genet. 5, 468.

Deatherage, D.E., Kepner, J.L., Bennett, A.F., Lenski, R.E., and Barrick, J.E. (2017). Specificity of genome evolution in experimental populations of *Escherichia coli* evolved at different temperatures. Proc. Natl. Acad. Sci. U. S. A. 114, E1904–E1912.

Desai, M.M., Walczak, A.M., and Fisher, D.S. (2012). Genetic diversity and the structure of genealogies in rapidly adapting populations. Genetics 193, 565–585.

Dinardo, S., Voelkel, K.A., Sternglanz, R., Reynolds, A.E., and Wright, A. (1982). *Escherichia coli* DNA topoisomerase I mutants have compensatory mutations in DNA gyrase genes. Cell 31, 43–51.

Ding, L., Ley, T.J., Larson, D.E., Miller, C. a., Koboldt, D.C., Welch, J.S., Ritchey, J.K., Young, M. a., Lamprecht, T., McLellan, M.D., et al. (2012). Clonal evolution in relapsed acute myeloid leukaemia revealed by whole-genome sequencing. Nature 481, 506–510.

Donovan, K.A., Atkinson, S.C., Kessans, S.A., Peng, F., Cooper, T.F., Griffin, M.D.W., Jameson, G.B., and Dobson, R.C.J. (2016). Grappling with anisotropic data, pseudo-merohedral twinning and pseudo-translational noncrystallographic symmetry: A case study involving pyruvate kinase. Acta Crystallogr. Sect. D Struct. Biol. 72, 512–519.

Fischer, S., Greipel, L., Klockgether, J., Dorda, M., Wiehlmann, L., Cramer, N., and Tümmler, B. (2017). Multilocus amplicon sequencing of *Pseudomonas aeruginosa* cystic fibrosis airways isolates collected prior to and after early antipseudomonal chemotherapy. J. Cyst. Fibros. 16, 346–352.

Fowler, D.M., and Fields, S. (2014). Deep mutational scanning: a new style of protein science. Nat. Methods 11, 801–807.

Frenkel, E.M., McDonald, M.J., Van Dyken, J.D., Kosheleva, K., Lang, G.I., and Desai, M.M. (2015). Crowded growth leads to the spontaneous evolution of semistable coexistence in laboratory yeast populations. Proc. Natl. Acad. Sci. 112, 11306–11311.

Furusawa, C., Horinouchi, T., and Maeda, T. (2018). Toward prediction and control of antibiotic-resistance evolution. Curr. Opin. Biotechnol. 54, 45–49.

Garst, A.D., Bassalo, M.C., Pines, G., Lynch, S.A., Halweg-Edwards, A.L., Liu, R., Liang, L., Wang, Z., Zeitoun, R., Alexander, W.G., et al. (2017). Genome-wide mapping of mutations at single-nucleotide resolution for protein, metabolic and genome engineering. Nat. Biotechnol. 35, 48–55.

Genovese, G., Kähler, A.K., Handsaker, R.E., Lindberg, J., Rose, S.A., Bakhoum, S.F., Chambert, K., Mick, E., Neale, B.M., Fromer, M., et al. (2014). Clonal hematopoiesis and blood-cancer risk inferred from blood DNA sequence. N. Engl. J. Med. 371, 2477–2487.

Gerrish, P.J., and Lenski, R.E. (1998). The fate of competing beneficial mutations in an asexual population. Genetica 102–103, 127–144.

Good, B.H., McDonald, M.J., Barrick, J.E., Lenski, R.E., and Desai, M.M. (2017). The dynamics of molecular evolution over 60,000 generations. Nature 551, 45–50.

Gresham, D., and Dunham, M.J. (2014). The enduring utility of continuous culturing in experimental evolution. Genomics 104, 399–405.

Hegreness, M., and Kishony, R. (2007). Analysis of genetic systems using experimental evolution and whole-genome sequencing. Genome Biol. 8, 201.

Hegreness, M., Shoresh, N., Hartl, D., and Kishony, R. (2006). An equivalence principle for the incorporation of favorable mutations in asexual populations. Science 311, 1615–1617.

Hong, J., Brandt, N., Abdul-Rahman, F., Yang, A., Hughes, T., and Gresham, D. (2018). An incoherent feedforward loop facilitates adaptive tuning of gene expression. Elife 7, 1–18.

Houdaigui, B. El, Forquet, R., Hindré, T., Schneider, D., Nasser, W., Reverchon, S., and Meyer, S. (2019). Bacterial genome architecture shapes global transcriptional regulation by DNA supercoiling. Nucleic Acids Res. 47, 5648–5657.

Jeong, H., Barbe, V., Lee, C.H., Vallenet, D., Yu, D.S., Choi, S.-H., Couloux, A., Lee, S.-W., Yoon, S.H., Cattolico, L., et al. (2009). Genome sequences of *Escherichia coli* B strains REL606 and BL21(DE3). J. Mol. Biol. 394, 644–652.

Khan, A.I., Dinh, D.M., Schneider, D., Lenski, R.E., and Cooper, T.F. (2011). Negative epistasis between beneficial mutations in an evolving bacterial population. Science 332, 1193–1196.

Kochanowski, K., Volkmer, B., Gerosa, L., Van Rijsewijk, B.R.H., Schmidt, A., and Heinemann, M. (2013). Functioning of a metabolic flux sensor in *Escherichia coli*. Proc. Natl. Acad. Sci. U. S. A. 110, 1130–1135.

Kryazhimskiy, S., Rice, D.P., Jerison, E.R., and Desai, M.M. (2014). Global epistasis makes adaptation predictable despite sequence-level stochasticity. Science 344, 1519–1522.

Kurnasov, O. V., Polanuyer, B.M., Ananta, S., Sloutsky, R., Tam, A., Gerdes, S.Y., and Osterman, A.L. (2003). Ribosylnicotinamide Kinase Domain of NadR Protein: Identification and Implications in NAD Biosynthesis. J. Bacteriol. 185, 698–698.

Landau, D.A., Carter, S.L., Stojanov, P., McKenna, A., Stevenson, K., Lawrence, M.S., Sougnez, C., Stewart, C., Sivachenko, A., Wang, L., et al. (2013). Evolution and impact of subclonal mutations in chronic lymphocytic leukemia. Cell 152, 714–726.

Lang, G.I., Rice, D.P., Hickman, M.J., Sodergren, E., Weinstock, G.M., Botstein, D., and Desai, M.M. (2013). Pervasive genetic hitchhiking and clonal interference in forty evolving yeast populations. Nature 500, 571–574.

Lee, S.Y., and Kim, H.U. (2015). Systems strategies for developing industrial microbial strains. Nat. Biotechnol. 33, 1061–1072.

Lenski, R.E., and Travisano, M. (1994). Dynamics of adaptation and diversification: a 10,000-generation experiment with bacterial populations. Proc. Natl. Acad. Sci. U. S. A. 91, 6808–6814.

Lenski, R.E., Rose, M.R., Simpson, S.C., and Tadler, S.C. (1991). Long-term experimental evolution in *Escherichia coli*. I. Adaptation and divergence during 2,000 generations. Am. Nat. 138, 1315–1341.

Levy, S.F., Blundell, J.R., Venkataram, S., Petrov, D.A., Fisher, D.S., and Sherlock, G. (2015). Quantitative evolutionary dynamics using high-resolution lineage tracking. Nature 519, 181–186.

Maddamsetti, R., Lenski, R.E., and Barrick, J.E. (2015). Adaptation, clonal interference, and frequency-dependent interactions in a long-term evolution experiment with *Escherichia coli*. Genetics 200, 619–631.

Marusyk, A., Almendro, V., and Polyak, K. (2012). Intra-tumour heterogeneity: A looking glass for cancer? Nat. Rev. Cancer 12, 323–334.

Marvig, R.L., Sommer, L.M., Molin, S., and Johansen, H.K. (2015). Convergent evolution and adaptation of *Pseudomonas aeruginosa* within patients with cystic fibrosis. Nat. Genet. 47, 57–64.

Massé, E., and Drolet, M. (1999). Relaxation of transcription-induced negative supercoiling is an essential function of *Escherichia coli* DNA topoisomerase I. J. Biol. Chem. 274, 16654–16658.

Mattevi, A., Valentini, G., Rizzi, M., Speranza, M.L., Bolognesi, M., and Coda, A. (1995). Crystal structure of *Escherichia coli* pyruvate kinase type I: molecular basis of the allosteric transition. Structure 3, 729–741.

McDonald, M.J. (2019). Microbial experimental evolution – a proving ground for evolutionary theory and a tool for discovery. EMBO Rep. 20, 1–14.

Merlo, L.M.F., Pepper, J.W., Reid, B.J., and Maley, C.C. (2006). Cancer as an evolutionary and ecological process. Nat. Rev. Cancer 6, 924–935.

Merlo, L.M.F., Shah, N.A., Li, X., Blount, P.L., Vaughan, T.L., Reid, B.J., and Maley, C.C. (2010). A comprehensive survey of clonal diversity measures in Barrett’s esophagus as biomarkers of progression to esophageal adenocarcinoma. Cancer Prev. Res. (Phila). 3, 1388–1397.

Newman, A.M., Bratman, S. V., To, J., Wynne, J.F., Eclov, N.C.W., Modlin, L.A., Liu, C.L., Neal, J.W., Wakelee, H.A., Merritt, R.E., et al. (2014). An ultrasensitive method for quantitating circulating tumor DNA with broad patient coverage. Nat. Med. 20, 548–554.

Newman, A.M., Lovejoy, A.F., Klass, D.M., Kurtz, D.M., Chabon, J.J., Scherer, F., Stehr, H., Liu, C.L., Bratman, S. V., Say, C., et al. (2016). Integrated digital error suppression for improved detection of circulating tumor DNA. Nat. Biotechnol. 34, 547–555.

Nielsen, J., and Keasling, J.D. (2016). Engineering cellular metabolism. Cell 164, 1185–1197.

Ostrowski, E.A., Woods, R.J., and Lenski, R.E. (2008). The genetic basis of parallel and divergent phenotypic responses in evolving populations of *Escherichia coli*. Proc. R. Soc. B 275, 277–284.

Park, S.-C., and Krug, J. (2007). Clonal interference in large populations. Proc. Natl. Acad. Sci. U. S. A. 104, 18135–18140.

Peng, F., Widmann, S., Wünsche, A., Duan, K., Donovan, K.A., Dobson, R.C.J., Lenski, R.E., and Cooper, T.F. (2018). Effects of beneficial mutations in *pykF* gene vary over time and across replicate populations in a long-term experiment with bacteria. Mol. Biol. Evol. 35, 202–210.

Perry, K., and Mondragón, A. (2003). Structure of a complex between *E. coli* DNA topoisomerase I and single-stranded DNA. Structure 11, 1349–1358.

Pruss, G.J., Manes, S.H., and Drlica, K. (1982). *Escherichia coli* DNA topoisomerase I mutants: Increased supercoiling is corrected by mutations near gyrase genes. Cell 31, 35–42.

R Core Team (2016). R: A Language and Environment for Statistical Computing (Vienna, Austria: R Foundation for Statistical Computing).

Raeside, C., Gaffé, J., Deatherage, D.E., Tenaillon, O., Briska, A.M., Ptashkin, R.N., Cruveiller, S., Médigue, C., Lenski, R.E., Barrick, J.E., et al. (2014). Large chromosomal rearrangements during a long-term evolution experiment with *Escherichia coli*. MBio 5, e01377–14.

Raffaelli, N., Lorenzi, T., Mariani, P.L., Emanuelli, M., Amici, A., Ruggieri, S., and Magni, G. (1999). The *Escherichia coli* NadR regulator is endowed with nicotinamide mononucleotide adenylyltransferase activity. J. Bacteriol. 181, 5509–5511.

Rainey, P.B., Remigi, P., Farr, A.D., and Lind, P.A. (2017). Darwin was right: where now for experimental evolution? Curr. Opin. Genet. Dev. 47, 102–109.

Renda, B.A., Hammerling, M.J., and Barrick, J.E. (2014). Engineering reduced evolutionary potential for synthetic biology. Mol. Biosyst. 10, 1668–1678.

Rugbjerg, P., and Sommer, M.O.A. (2019). Overcoming genetic heterogeneity in industrial fermentations. Nat. Biotechnol. 37, 869–876.

Rugbjerg, P., Myling-Petersen, N., Porse, A., Sarup-Lytzen, K., and Sommer, M.O.A. (2018). Diverse genetic error modes constrain large-scale bio-based production. Nat. Commun. 9, 787.

Ryall, B., Eydallin, G., and Ferenci, T. (2012). Culture history and population heterogeneity as determinants of bacterial adaptation: the adaptomics of a single environmental transition. Microbiol. Mol. Biol. Rev. 76, 597–625.

Sandoval, C.M., Ayson, M., Moss, N., Lieu, B., Jackson, P., Gaucher, S.P., Horning, T., Dahl, R.H., Denery, J.R., Abbott, D.A., et al. (2014). Use of pantothenate as a metabolic switch increases the genetic stability of farnesene producing *Saccharomyces cerevisiae*. Metab. Eng. 25, 215–226.

Schmitt, M.W., Kennedy, S.R., Salk, J.J., Fox, E.J., Hiatt, J.B., and Loeb, L.A. (2012). Detection of ultra-rare mutations by next-generation sequencing. Proc. Natl. Acad. Sci. U. S. A. 109, 14508–14513.

Siddiquee, K.A.Z., Arauzo-Bravo, M.J., and Shimizu, K. (2004). Effect of a pyruvate kinase (*pykF-gene*) knockout mutation on the control of gene expression and metabolic fluxes in *Escherichia coli*. FEMS Microbiol. Lett. 235, 25–33.

Stefani, S., Campana, S., Cariani, L., Carnovale, V., Colombo, C., Lleo, M.M., Iula, V.D., Minicucci, L., Morelli, P., Pizzamiglio, G., et al. (2017). Relevance of multidrug-resistant *Pseudomonas aeruginosa* infections in cystic fibrosis. Int. J. Med. Microbiol. 307, 353–362.

Tan, K., Zhou, Q., Cheng, B., Zhang, Z., Joachimiak, A., and Tse-Dinh, Y.C. (2015). Structural basis for suppression of hypernegative DNA supercoiling by *E. coli* topoisomerase I. Nucleic Acids Res. 43, 11031–11046.

Tenaillon, O., Rodríguez-Verdugo, A., Gaut, R.L., McDonald, P., Bennett, A.F., Long, A.D., and Gaut, B.S. (2012). The molecular diversity of adaptive convergence. Science 335, 457–461.

Tenaillon, O., Barrick, J.E., Ribeck, N., Deatherage, D.E., Blanchard, J.L., Dasgupta, A., Wu, G.C., Wielgoss, S., Cruveiller, S., Médigue, C., et al. (2016). Tempo and mode of genome evolution in a 50,000-generation experiment. Nature 536, 165–170.

Thomas, R.K., Nickerson, E., Simons, J.F., Jänne, P.A., Tengs, T., Yuza, Y., Garraway, L.A., LaFramboise, T., Lee, J.C., Shah, K., et al. (2006). Sensitive mutation detection in heterogeneous cancer specimens by massively parallel picoliter reactor sequencing. Nat. Med. 12, 852–855.

Traverse, C.C., Mayo-Smith, L.M., Poltak, S.R., and Cooper, V.S. (2013). Tangled bank of experimentally evolved *Burkholderia* biofilms reflects selection during chronic infections. Proc. Natl. Acad. Sci. U. S. A. 110, E250–E259.

Venkataram, S., Dunn, B., Li, Y., Agarwala, A., Chang, J., Ebel, E.R., Geiler-Samerotte, K., Hérissant, L., Blundell, J.R., Levy, S.F., et al. (2016). Development of a comprehensive genotype-to-fitness map of adaptation-driving mutations in yeast. Cell 166, 1585–1596.E22.

Watson, C.J., Papula, A.L., Poon, G.Y.P., Wong, W.H., Young, A.L., Druley, T.E., Fisher, D.S., and Blundell, J.R. (2020). The evolutionary dynamics and fitness landscape of clonal hematopoiesis. Science 367, 1449–1454.

Wei, X., and Zhang, J. (2019). Patterns and mechanisms of diminishing returns from beneficial mutations. Mol. Biol. Evol. 36, 1008–1021.

Wickham, H. (2016). ggplot2: Elegant Graphics for Data Analysis (New York: Springer-Verlag).

Winstanley, C., O’Brien, S., and Brockhurst, M.A. (2016). *Pseudomonas aeruginosa* evolutionary adaptation and diversification in cystic fibrosis chronic lung infections. Trends Microbiol. 24, 327–337.

Wiser, M.J., Ribeck, N., and Lenski, R.E. (2013). Long-term dynamics of adaptation in asexual populations. Science 342, 1364–1367.

Woods, R., Schneider, D., Winkworth, C.L., Riley, M.A., and Lenski, R.E. (2006). Tests of parallel molecular evolution in a long-term experiment with *Escherichia coli*. Proc. Natl. Acad. Sci. U. S. A. 103, 9107–9712.

Woods, R.J., Barrick, J.E., Cooper, T.F., Shrestha, U., Kauth, M.R., and Lenski, R.E. (2011). Second-order selection for evolvability in a large *Escherichia coli* population. Science 331, 1433–1436.

Zelder, O., and Hauer, B. (2000). Environmentally directed mutations and their impact on industrial biotransformation and fermentation processes. Curr. Opin. Microbiol. 3, 248–251.

