## Supplemental Information for "Profiling the initial burst of beneficial genetic diversity in clonal cell populations to anticipate evolution"

**Table S1.** Adapter sequences

| Name | Sequence |
|---|---|
| UTBC52 | /5Phos/ACTG**NNNNNNNNNNNNAGATCGGAAGAGC**ACACGTCTGAACTCCAGTCAC**AACATA**ATCTCGTATGCCGTCTTCTCGTTG |
| UTBC75 | /5Phos/ACTG**NNNNNNNNNNNNAGATCGGAAGAGC**ACACGTCTGAACTCCAGTCAC**ACGCCG**ATCTCGTATGCCGTCTTCTCGTTG |
| TSBC25 | /5Phos/ACTG**NNNNNNNNNNNNAGATCGGAAGAGC**ACACGTCTGAACTCCAGTCAC**ACTGAT**ATCTCGTATGCCGTCTTCTCGTTG |
| TSBC14 | /5Phos/ACTG**NNNNNNNNNNNNAGATCGGAAGAGC**ACACGTCTGAACTCCAGTCAC**AGTTCC**ATCTCGTATGCCGTCTTCTCGTTG |
| UTBC64 | /5Phos/ACTG**NNNNNNNNNNNNAGATCGGAAGAGC**ACACGTCTGAACTCCAGTCAC**ATACGA**ATCTCGTATGCCGTCTTCTCGTTG |
| TSBC34 | /5Phos/ACTG**NNNNNNNNNNNNAGATCGGAAGAGC**ACACGTCTGAACTCCAGTCAC**CATGGC**ATCTCGTATGCCGTCTTCTCGTTG |
| TSBC36 | /5Phos/ACTG**NNNNNNNNNNNNAGATCGGAAGAGC**ACACGTCTGAACTCCAGTCAC**CCAACA**ATCTCGTATGCCGTCTTCTCGTTG |
| UTBC63 | /5Phos/ACTG**NNNNNNNNNNNNAGATCGGAAGAGC**ACACGTCTGAACTCCAGTCAC**CCCCAC**ATCTCGTATGCCGTCTTCTCGTTG |
| TSBC37 | /5Phos/ACTG**NNNNNNNNNNNNAGATCGGAAGAGC**ACACGTCTGAACTCCAGTCAC**CGGAAT**ATCTCGTATGCCGTCTTCTCGTTG |
| UTBC58 | /5Phos/ACTG**NNNNNNNNNNNNAGATCGGAAGAGC**ACACGTCTGAACTCCAGTCAC**CGGTTA**ATCTCGTATGCCGTCTTCTCGTTG |
| TSBC23 | /5Phos/ACTG**NNNNNNNNNNNNAGATCGGAAGAGC**ACACGTCTGAACTCCAGTCAC**GAGTGG**ATCTCGTATGCCGTCTTCTCGTTG |
| TSBC09 | /5Phos/ACTG**NNNNNNNNNNNNAGATCGGAAGAGC**ACACGTCTGAACTCCAGTCAC**GATCAG**ATCTCGTATGCCGTCTTCTCGTTG |
| UTBC86 | /5Phos/ACTG**NNNNNNNNNNNNAGATCGGAAGAGC**ACACGTCTGAACTCCAGTCAC**GGCGGT**ATCTCGTATGCCGTCTTCTCGTTG |
| TSBC20 | /5Phos/ACTG**NNNNNNNNNNNNAGATCGGAAGAGC**ACACGTCTGAACTCCAGTCAC**GTGGCC**ATCTCGTATGCCGTCTTCTCGTTG |
| UTBC69 | /5Phos/ACTG**NNNNNNNNNNNNAGATCGGAAGAGC**ACACGTCTGAACTCCAGTCAC**GTTATT**ATCTCGTATGCCGTCTTCTCGTTG |
| UTBC56 | /5Phos/ACTG**NNNNNNNNNNNNAGATCGGAAGAGC**ACACGTCTGAACTCCAGTCAC**TAAGAA**ATCTCGTATGCCGTCTTCTCGTTG |
| TSBC10 | /5Phos/ACTG**NNNNNNNNNNNNAGATCGGAAGAGC**ACACGTCTGAACTCCAGTCAC**TAGCTT**ATCTCGTATGCCGTCTTCTCGTTG |
| TSBC45 | /5Phos/ACTG**NNNNNNNNNNNNAGATCGGAAGAGC**ACACGTCTGAACTCCAGTCAC**TCATTC**ATCTCGTATGCCGTCTTCTCGTTG |
| UTBC51 | /5Phos/ACTG**NNNNNNNNNNNNAGATCGGAAGAGC**ACACGTCTGAACTCCAGTCAC**TGAAGG**ATCTCGTATGCCGTCTTCTCGTTG |
| UTBC94 | /5Phos/ACTG**NNNNNNNNNNNNAGATCGGAAGAGC**ACACGTCTGAACTCCAGTCAC**TTCTAT**ATCTCGTATGCCGTCTTCTCGTTG |
| Univ-F | AATGATACGGCGACCACCGAGATCTACACTCTTTCCCTACACGAC**GCTCTTCCGATCT** |

/5Phos/ indicates that an oligonucleotide was chemically synthesized with a 5′ monophosphate. **Red bases** are sample barcodes. **Blue bases** are unique molecular identifiers (UMIs) that were synthesized with a mixture of all four bases at each N position. **Green bases** show where each sequence was individually annealed to Univ-F through the sequence and extended to make the double-stranded end of the adaptor that includes the unique molecular identifier before it was ligated to the DNA sample.

**Table S2.** Probe sequences

| Name | Sequence | Start | End |
|---|---|---|---|
| ybaL_1 | TATTTTTCATATTTTACATCCGGCAACCACCGTTTACCCCGTCACCACCTCACCCGCCGG | 473596 | 473655 |
| ybaL_2 | TGGCGTTTCCAGCAGTTCCAGCATGGTACGGGCGATTTCACGCTCGCCCATCACTACCTG | 473656 | 473715 |
| ybaL_3 | ATTCGCACCACGTTCGGTGATATACGCCACTTCATCGTCATAATGGGCGCGGGCAATAAT | 473716 | 473775 |
| ybaL_4 | CTCAATATCCGGATTTTTCGCGCGGGCAGATGCCACAATCTCACCCGCTTCATAACCGTT | 473776 | 473835 |
| ybaL_5 | GGGAATCGTCAGGATCAGCCATTTTGCACATTCCAGATGCGCCAGTTGCATAATTTCTTC | 473836 | 473895 |
| ybaL_6 | GTTCGCCGCATTGCCCAATACTGCGCGGACCCCGCGCTCTCGCAGCTCATCAACACGGGT | 473896 | 473955 |
| ybaL_7 | TCGTGACGTCTCAATCACCACCAGCGGAATATCAGAGGCGAGCAATTTCTCCCCCAGCAG | 473956 | 474015 |
| ybaL_8 | GCTGCCTACACGACCGTAACCCACCAGTAGCGCATGGTTGCAAATATCCACTGGGATCTG | 474016 | 474075 |
| ybaL_9 | CTTCTCTTCTTCGATTGCCTCTTCCAGCGTCTGCTCTTCCAGCGTTTCGGTCTTCGCCAG | 474076 | 474135 |
| ybaL_10 | ATATTTCTCCAGTAGTGCGAACAGTACCGGGTTGAGCATAATCGACAGGATCGCCCCTGC | 474136 | 474195 |
| ybaL_11 | CAGTACCAGGTTTTGTCCGGCCTGCGGCAGTAAATTCAATGCCATTCCCAGTCCCGCCAG | 474196 | 474255 |
| ybaL_12 | GATAAACGCGAACTCACCAATCTGCGCCAGGCTGGCGGCGATGGTTAATGCCGTACGTTG | 474256 | 474315 |
| ybaL_13 | GGAGTGACCAAACAGTCGCACCAGGAAAAATGCGGCTAACGACTTACCAAACAGAATAAT | 474316 | 474375 |
| ybaL_14 | CGCCAGCGTCGCCAGCACTGCCAGCGGTTGCTGAATCAGAATTAACGGATCAAACAACAT | 474376 | 474435 |
| ybaL_15 | CCCGACGGAGACAAAAAACAGCACCGCAAACGCGTCGCGCAATGGCAGCGTATCGTGGGC | 474436 | 474495 |
| ybaL_16 | GGCACGGTGACTCAGTTCAGACTCGTTCAGTACCATCCCGGCAAAGAACGCACCGAGTGC | 474496 | 474555 |
| ybaL_17 | AAAGGAGACATCAAACAGCTCTACCGCACCAAAGGCAACCCCTAACGCCAGCGCCAGCAC | 474556 | 474615 |
| ybaL_18 | CGACAGGGTAAACAGCTCGCGAGAACCGGTTGCCGCGCTGCGTGCCATAATCCACGGCAC | 474616 | 474675 |
| ybaL_19 | CAGACGGCGACCTACCAGCATCATAATGGCGATAAATGCGATCACTTTGCCGATGGTGAT | 474676 | 474735 |
| ybaL_20 | CCCCATATCGACTGCAAGAGTGGCAAAGCCCACATCGCCCTGTTCCATCATTCCTGCCAC | 474736 | 474795 |
| ybaL_21 | TGCGGGCAGCAACACCAGCGTCAGAACCATTACCAGGTCTTCCACAATCAACCAACCGAT | 474796 | 474855 |
| ybaL_22 | GGCGATTTGCCCACGCTGACTGTCAATTAATTGCCGTTCTTCAAGTGCGCGCAGTAACAC | 474856 | 474915 |
| ybaL_23 | CACGGTACTGGCGGTGGAAAGACATAAACCGAACACGATACCGGTCATTAACGACCAGCC | 474916 | 474975 |
| ybaL_24 | CAGCACGGCAGAGAGCGCCATACCCAGCAGCGTCGCCACGGCTATCTGGGCGATCGCACC | 474976 | 475035 |
| ybaL_25 | GGGAATGGCGATGGCCTTTACCGCCATCAAATCCTTCAGCGAAAAGTGCAAACCGACGCC | 475036 | 475095 |
| ybaL_26 | AAACATCAACAGAATGACGCCCAGTTCAGCCAGTTCCGGGGCAAGCTTGGTATCGGCAAC | 475096 | 475155 |
| ybaL_27 | AAAGCCCGGAGTGAATGGTCCTGCCAGCACACCCGCTAACAGATATCCCACCAGAGGAGA | 475156 | 475215 |
| ybaL_28 | AATACGTAGTTTATTGGCCAGCATGCCGAGGATAAAGGCGAGCACAAGGCCGCCAACAAT | 475216 | 475275 |
| ybaL_29 | GGTGGTGATAAGCGGGGTGGCGTGATGCATTCCGTCTCCTTTTCCTGGTGGTTATTGTCC | 475276 | 475335 |
| ybaL_30 | ATTTTTGGCCGGGAAAACCAAAATTACAGGTAATAGTTTATGACAATTTCATTGATGATG | 475336 | 475395 |
| ybaL_31 | TTCATGAATAATTATTGAATTTTGCAGAAAAATGGAATTAGCTGCAAAAAAAGCACGGAT | 475396 | 475455 |
| topA_1 | GCCTTACTGGCAACTTTGGATTTTGCATGCTAATAAAGTTGCGTATCGGATTTTATCAGG | 1329119 | 1329178 |
| topA_2 | TACAGTGTGACGCTTTCGTCAATCTGGCAATAGATTTGCTTGACATTCGACCAAAATTCC | 1329179 | 1329238 |
| topA_3 | GTCGTGCTATAGCGCCTGTAGGCCAAGACCTGTTAACTCAGTCACCTGAATTTTCGTGAA | 1329239 | 1329298 |
| topA_4 | CAGAGTCACGACAAGGGGTTGATATCCGCAGAGAGCGAGTCCATATCGGTAACTCGTTGC | 1329299 | 1329358 |
| topA_5 | CAGTGGAAGGTTTATCAACGTGCGACGCATTCCTGGAAGAATCAAATTAGGTAAGGTGAA | 1329359 | 1329418 |
| topA_6 | TATGGGTAAAGCTCTTGTCATCGTTGAGTCCCCGGCAAAAGCCAAAACGATCAACAAGTA | 1329419 | 1329478 |
| topA_7 | TCTGGGTAGTGACTACGTGGTGAAATCCAGCGTCGGTCACATCCGCGATTTGCCGACCAG | 1329479 | 1329538 |
| topA_8 | TGGCTCAGCTGCCAAAAAGAGTGCCGACTCTACCTCCACCAAGACGGCTAAAAAGCCTAA | 1329539 | 1329598 |
| topA_9 | AAAGGATGAACGTGGCGCTCTCGTCAACCGTATGGGGGTTGACCCGTGGCACAATTGGGA | 1329599 | 1329658 |
| topA_10 | GGCGCACTATGAAGTGTTGCCTGGTAAAGAGAAGGTCGTCTCTGAACTGAAACAACTGGC | 1329659 | 1329718 |
| topA_11 | TGAAAAAGCCGACCACATCTATCTCGCAACCGACCTTGACCGCGAAGGGGAAGCCATTGC | 1329719 | 1329778 |
| topA_12 | ATGGCACCTGCGGGAAGTGATTGGGGGTGATGATGCGCGCTATAGCCGAGTGGTGTTTAA | 1329779 | 1329838 |
| topA_13 | CGAAATTACTAAAAACGCGATCCGCCAGGCATTTAACAAACCGGGTGAGCTGAATATTGA | 1329839 | 1329898 |
| topA_14 | TCGTGTTAATGCCCAGCAGGCGCGTCGCTTTATGGACCGCGTGGTGGGGTATATGGTTTC | 1329899 | 1329958 |
| topA_15 | GCCGCTGCTATGGAAAAAGATCGCTCGTGGTCTGTCTGCCGGTCGTGTGCAGTCGGTGGC | 1329959 | 1330018 |
| topA_16 | GGTTCGCCTGGTGGTCGAGCGTGAGCGTGAAATTAAAGCGTTCGTGCCGGAAGAGTTCTG | 1330019 | 1330078 |
| topA_17 | GGAAGTCGATGCCAGCACGACCACGCCATCTGGTGAAGCGTTGGCGTTACAGGTGACTCA | 1330079 | 1330138 |
| topA_18 | TCAGAACGACAAACCGTTCCGTCCGGTCAACAAAGAACAAACTCAGGCTGCGGTAAGTCT | 1330139 | 1330198 |

| Name | Sequence | Start | End |
|---|---|---|---|
| topA_19 | GCTGGAAAAAGCGCGCTACAGCGTGCTGGAACGTGAAGACAAACCGACAACCAGTAAACC | 1330199 | 1330258 |
| topA_20 | TGGCGCTCCTTTTATTACCTCTACGCTGCAACAAGCTGCCAGCACCCGTCTTGGATTTGG | 1330259 | 1330318 |
| topA_21 | CGTGAAAAAAACCATGATGATGGCGCAGCGTTTGTATGAAGCAGGCTATATCACTTACAT | 1330319 | 1330378 |
| topA_22 | GCGTACCGACTCCACTAACCTGAGTCAGGACGCGGTAAATATGGTTCGCGGTTATATCAG | 1330379 | 1330438 |
| topA_23 | CGATAATTTTGGTAAGAAATATCTGCCGGAAAGTCCGAATCAGTACGCCAGCAAAGAAAA | 1330439 | 1330498 |
| topA_24 | CTCACAGGAAGCGCACGAAGCGATTCGTCCTTCTGACGTCAATGTGATGGCGGAATCGCT | 1330499 | 1330558 |
| topA_25 | GAAGGATATGGAAGCAGATGCGCAGAAACTGTACCAGTTAATCTGGCGTCAGTTCGTTGC | 1330559 | 1330618 |
| topA_26 | CTGCCAGATGACCCCAGCGAAATATGACTCCACGACGCTGACCGTTGGTGCGGGCGATTT | 1330619 | 1330678 |
| topA_27 | CCGCCTGAAAGCACGCGGTCGTATTTTGCGTTTTGATGGCTGGACAAAAGTGATGCCTGC | 1330679 | 1330738 |
| topA_28 | GTTGCGTAAAGGCGATGAAGATCGCATCTTACCAGCAGTTAATAAAGGCGATGCTCTGAC | 1330739 | 1330798 |
| topA_29 | GCTCGTTGAACTTACACCAGCCCAGCACTTTACCAAGCCGCCAGCCCGTTTCAGTGAAGC | 1330799 | 1330858 |
| topA_30 | ATCGCTGGTTAAAGAGCTGGAAAAACGCGGTATCGGTCGTCCGTCTACCTATGCGTCGAT | 1330859 | 1330918 |
| topA_31 | CATTTCGACCATTCAGGATCGTGGCTACGTGCGAGTAGAAAATCGTCGTTTCTATGCGGA | 1330919 | 1330978 |
| topA_32 | AAAAATGGGCGAAATCGTCACCGATCGCCTTGAAGAAAATTTCCGCGAGTTAATGAACTA | 1330979 | 1331038 |
| topA_33 | CGATTTTACCGCGCAGATGGAAAACAGCCTCGACCAGGTGGCAAATCACGAAGCAGAGTG | 1331039 | 1331098 |
| topA_34 | GAAAGCTGTACTGGATCACTTCTTCTCGGATTTCACCCAGCAGTTAGATAAAGCTGAAAA | 1331099 | 1331158 |
| topA_35 | AGATCCGGAAGAGGGTGGTATGCGCCCGAACCAGATGGTTCTGACCAGCATTGACTGCCC | 1331159 | 1331218 |
| topA_36 | GACTTGTGGTCGCAAAATGGGGATTCGCACAGCGAGCACCGGGGTATTCCTTGGCTGTTC | 1331219 | 1331278 |
| topA_37 | TGGCTATGCGCTGCCGCCGAAAGAGCGTTGCAAAACCACCATTAACCTGGTGCCGGAAAA | 1331279 | 1331338 |
| topA_38 | CGAAGTGCTGAACGTGCTGGAAGGCGAAGATGCTGAAACCAACGCGCTGCGCGCAAAACG | 1331339 | 1331398 |
| topA_39 | TCGTTGCCCGAAATGCGGCACGGCGATGGACAGCTATCTCATCGATCCGAAACGTAAGTT | 1331399 | 1331458 |
| topA_40 | GCATGTCTGTGGTAATAACCCAACCTGCGACGGTTACGAGATCGAAGAGGGCGAATTCCG | 1331459 | 1331518 |
| topA_41 | CATTAAAGGTTATGACGGCCCGATCGTTGAGTGTGAAAAATGTGGCTCTGAAATGCACCT | 1331519 | 1331578 |
| topA_42 | GAAAATGGGGCGTTTCGGTAAATACATGGCCTGCACCAACGAAGAGTGTAAAAACACACG | 1331579 | 1331638 |
| topA_43 | TAAGATTTTACGTAACGGCGAAGTGGCACCACCGAAAGAAGATCCGGTGCCATTACCTGA | 1331639 | 1331698 |
| topA_44 | GCTGCCGTGCGAAAAATCAGATGCTTATTTCGTGCTGCGTGACGGTGCTGCCGGTGTGTT | 1331699 | 1331758 |
| topA_45 | CCTGGCTGCCAACACTTTCCCGAAATCGCGTGAAACGCGTGCGCCACTGGTGGAAGAGCT | 1331759 | 1331818 |
| topA_46 | TTATCGCTTCCGCGACCGTCTGCCGGAAAAACTGCGTTATCTGGCCGATGCGCCACAGCA | 1331819 | 1331878 |
| topA_47 | GGATCCGGAAGGTAATAAGACCATGGTTCGCTTTAGCCGTAAAACCAAACAGCAATATGT | 1331879 | 1331938 |
| topA_48 | CTCTTCGGAAAAAGACGGAAAGGCGACTGGCTGGTCAGCATTTTATGTTGATGGCAAATG | 1331939 | 1331998 |
| topA_49 | GGTTGAAGGAAAAAAATAACCTTTAATTCTGTCAGGTTTTTATAAACAAAGGGTCGCGAA | 1331999 | 1332058 |
| pykF_1 | AACGCTGTTTTTGTTTTCCTTTTGGATTAATTTCAGCGTATAATGCGCGCCAATTGACTC | 1732685 | 1732744 |
| pykF_2 | TTGAATGGTTTCAGCACTTTGGACTGTAGAACTCAACGACTCAAAAACAGGCACTCACGT | 1732745 | 1732804 |
| pykF_3 | TGGGCTGAGACACAAGCACACATTCCTCTGCACGCTTTTTCGATGTCACCTATCCTTAGA | 1732805 | 1732864 |
| pykF_4 | GCGAGGCACCACCACTTTCGTAATACCGGATTCGCTTTCCGGCAGTGCGCCCAGAAAGCA | 1732865 | 1732924 |
| pykF_5 | AGTTTCTCCCATCCTTCTCAACTTAAAGACTAAGACTGTCATGAAAAAGACCAAAATTGT | 1732925 | 1732984 |
| pykF_6 | TTGCACCATCGGACCGAAAACCGAATCTGAAGAGATGTTAGCTAAAATGCTGGACGCTGG | 1732985 | 1733044 |
| pykF_7 | CATGAACGTTATGCGTCTGAACTTCTCTCATGGTGACTATGCAGAACACGGTCAGCGCAT | 1733045 | 1733104 |
| pykF_8 | TCAGAATCTGCGCAACGTGATGAGCAAAACTGGTAAAACCGCCGCTATCCTGCTTGATAC | 1733105 | 1733164 |
| pykF_9 | CAAAGGTCCGGAAATCCGCACCATGAAACTGGAAGGCGGTAACGACGTTTCTCTGAAAGC | 1733165 | 1733224 |
| pykF_10 | TGGTCAGACCTTTACTTTCACCACTGATAAATCTGTTATCGGCAACAGCGAAATGGTTGC | 1733225 | 1733284 |
| pykF_11 | GGTAACGTATGAAGGTTTCACTACTGACCTGTCTGTTGGCAACACCGTACTGGTTGACGA | 1733285 | 1733344 |
| pykF_12 | TGGTCTGATCGGTATGGAAGTTACCGCCATTGAAGGTAACAAAGTTATCTGTAAAGTGCT | 1733345 | 1733404 |
| pykF_13 | GAACAACGGTGACCTGGGCGAAAACAAAGGTGTGAACCTGCCTGGCGTTTCCATTGCTCT | 1733405 | 1733464 |
| pykF_14 | GCCAGCACTGGCTGAAAAAGACAAACAGGACCTGATCTTTGGTTGCGAACAAGGCGTAGA | 1733465 | 1733524 |
| pykF_15 | CTTTGTTGCTGCTTCCTTTATTCGTAAGCGTTCTGACGTTATCGAAATCCGTGAGCACCT | 1733525 | 1733584 |
| pykF_16 | GAAAGCGCACGGCGGCGAAAACATCCACATCATCTCCAAAATCGAAAACCAGGAAGGCCT | 1733585 | 1733644 |
| pykF_17 | CAACAACTTCGACGAAATCCTCGAAGCCTCTGACGGCATCATGGTTGCGCGTGGCGACCT | 1733645 | 1733704 |
| pykF_18 | GGGTGTAGAAATCCCGGTAGAAGAAGTTATCTTCGCCCAGAAGATGATGATCGAAAAATG | 1733705 | 1733764 |
| pykF_19 | TATCCGTGCACGTAAAGTCGTTATCACTGCGACCCAGATGCTGGATTCCATGATCAAAAA | 1733765 | 1733824 |
| pykF_20 | CCCACGCCCGACTCGCGCAGAAGCCGGTGACGTTGCAAACGCCATCCTCGACGGTACTGA | 1733825 | 1733884 |

| Name | Sequence | Start | End |
|---|---|---|---|
| pykF_21 | CGCAGTGATGCTGTCTGGTGAATCCGCAAAAGGTAAATACCCGCTGGAAGCGGTTTCTAT | 1733885 | 1733944 |
| pykF_22 | CATGGCGACCATCTGCGAACGTACCGACCGCGTGATGAACAGCCGTCTCGAGTTCAACAA | 1733945 | 1734004 |
| pykF_23 | TGACAACCGTAAACTGCGCATTACCGAAGCGGTATGCCGTGGTGCCGTTGAAACTGCTGA | 1734005 | 1734064 |
| pykF_24 | AAAACTGGATGCTCCGCTGATCGTGGTTGCTACTCAGGGCGGTAAATCTGCTCGCGCAGT | 1734065 | 1734124 |
| pykF_25 | ACGTAAATACTTCCCGGATGCCACCATCCTGGCACTGACCACCAACGAAAAAACGGCTCA | 1734125 | 1734184 |
| pykF_26 | TCAGTTGGTACTGAGCAAAGGCGTTGTGCCGCAGCTTGTTAAAGAGATCACTTCTACTGA | 1734185 | 1734244 |
| pykF_27 | TGATTTCTACCGTCTGGGTAAAGAACTGGCTCTGCAGAGCGGTCTGGCACACAAAGGTGA | 1734245 | 1734304 |
| pykF_28 | CGTTGTAGTTATGGTTTCTGGTGCACTGGTACCGAGCGGCACTACTAACACCGCATCTGT | 1734305 | 1734364 |
| pykF_29 | TCACGTCCTGTAATATTGCTTTTGTGAATTAATTTGTATATCGAAGCGCCCTGATGGGCG | 1734365 | 1734424 |
| spoT_1 | GCAAATTGTTGGCAGACTGAACCTGATTTCAGTATCATGCCCAGTCATTTCTTCACCTGT | 3760389 | 3760448 |
| spoT_2 | GGAGCTTTTTAAGTATGGCACGCGTAACTGTTCAGGACGCTGTAGAGAAAATTGGTAACC | 3760449 | 3760508 |
| spoT_3 | GTTTTGACCTGGTACTGGTCGCCGCGCGTCGCGCTCGTCAGATGCAGGTAGGCGGAAAGG | 3760509 | 3760568 |
| spoT_4 | ATCCGCTGGTACCGGAAGAAAACGATAAAACCACTGTAATCGCGCTGCGCGAAATCGAAG | 3760569 | 3760628 |
| spoT_5 | AAGGTCTGATCAACAACCAGATCCTCGACGTTCGCGAACGCCAGGAACAGCAAGAGCAGG | 3760629 | 3760688 |
| spoT_6 | AAGCCGCTGAATTACAAGCCGTTACCGCTATTGCTGAAGGTCGTCGTTAATCACAAAGCG | 3760689 | 3760748 |
| spoT_7 | GGTCGCCCTTGTATCTGTTTGAAAGCCTGAATCAACTGATTCAAACCTACCTGCCGGAAG | 3760749 | 3760808 |
| spoT_8 | ACCAAATCAAGCGTCTGCGGCAGGCGTATCTCGTTGCACGTGATGCTCACGAGGGGCAAA | 3760809 | 3760868 |
| spoT_9 | CACGTTCAAGCGGTGAACCCTATATCACGCACCCGGTAGCGGTTGCCTGCATTCTGGCCG | 3760869 | 3760928 |
| spoT_10 | AGATGAAACTCGACTATGAAACGCTGATGGCGGCGCTGCTGCATGACGTGATTGAAGATA | 3760929 | 3760988 |
| spoT_11 | CTCCCGCCACCTACCAGGATATGGAACAGCTTTTTGGTAAAAGCGTCGCCGAGCTGGTAG | 3760989 | 3761048 |
| spoT_12 | AGGGGGTGTCGAAACTTGATAAACTCAAGTTCCGCGATAAGAAAGAGGCGCAGGCCGAAA | 3761049 | 3761108 |
| spoT_13 | ACTTTCGCAAGATGATTATGGCGATGGTGCAGGATATCCGCGTCATCCTCATCAAACTTG | 3761109 | 3761168 |
| spoT_14 | CCGACCGTACCCACAACATGCGCACGCTGGGCTCACTTCGCCCGGACAAACGTCGCCGCA | 3761169 | 3761228 |
| spoT_15 | TCGCCCGTGAAACTCTCGAAATTTACAGCCCGCTGGCGCACCGTTTAGGTATCCACCACA | 3761229 | 3761288 |
| spoT_16 | TTAAAACCGAACTCGAAGAGCTGGGTTTTGAGGCGCTGTATCCCAATCGTTACCGCGTAA | 3761289 | 3761348 |
| spoT_17 | TTAAAGAAGTGGTGAAAGCCGCGCGCGGCAACCGTAAAGAGATGATCCAAAAAATCCTCT | 3761349 | 3761408 |
| spoT_18 | CTGAAATCGAAGGGCGTTTGCAGGAAGCGGGAATACCGTGCCGCGTCAGTGGTCGCGAAA | 3761409 | 3761468 |
| spoT_19 | AGCATCTTTATTCGATTTACTGCAAAATGGTGCTCAAAGAGCAGCGTTTTCACTCAATCA | 3761469 | 3761528 |
| spoT_20 | TGGACATCTACGCTTTCCGCGTGATCGTCAATGATTCTGACACCTGTTATCGCGTGCTGG | 3761529 | 3761588 |
| spoT_21 | GCCAGATGCACAGCCTGTACAAGCCGCGTCCGGGCCGCGTGAAAGACTATATCGCCATTC | 3761589 | 3761648 |
| spoT_22 | CAAAAGCGAACGGCTATCAGTCGTTGCACACCTCGATGATTGGCCCGCACAGCGTGCCGG | 3761649 | 3761708 |
| spoT_23 | TTGAGGTCCAGATCCGTACCGAAGATATGGATCAGATGGCGGAGATGGGTGTTGCCGCGC | 3761709 | 3761768 |
| spoT_24 | ACTGGGCTTATAAAGAGCACGGCGAAACCAGTACTACCGCACAAATCCGCGCCCAGCGCT | 3761769 | 3761828 |
| spoT_25 | GGATGCAAAGCCTGCTGGAGCTGCAACAGAGCGCCGGTAGTTCGTTTGAATTTATCGAGA | 3761829 | 3761888 |
| spoT_26 | GCGTTAAATCCGATCTCTTCCCGGATGAGATTTACGTTTTCACACCGGAAGGGCGCATTG | 3761889 | 3761948 |
| spoT_27 | TCGAGCTGCCTGCCGGTGCAACGCCCGTCGACTTCGCTTATGCAGTGCATACCGATATCG | 3761949 | 3762008 |
| spoT_28 | GTCATGCCTGCGTGGGCGCACGCGTTGACCGCCAGCCTTACCCGCTGTCGCAGCCGCTTA | 3762009 | 3762068 |
| spoT_29 | CCAGCGGTCAAACCGTTGAAATCATTACCGCTCCGGGCGCTCGCCCGAATGCCGCTTGGC | 3762069 | 3762128 |
| spoT_30 | TGAACTTTGTCGTTAGCTCGAAAGCGCGCGCCAAAATTCGTCAGTTGCTGAAAAACCTCA | 3762129 | 3762188 |
| spoT_31 | AGCGTGATGATTCTGTAAGCCTGGGCCGTCGTCTGCTCAACCATGCTTTGGGTGGTAGCC | 3762189 | 3762248 |
| spoT_32 | GTAAGCTGAATGAAATCCCGCAGGAAAATATTCAGCGCGAGCTGGATCGCATGAAGCTGG | 3762249 | 3762308 |
| spoT_33 | CAACGCTTGACGATCTGCTGGCAGAAATCGGACTTGGTAACGCAATGAGCGTGGTGGTCG | 3762309 | 3762368 |
| spoT_34 | CGAAAAATCTGCAACATGGGGACGCCTCCATTCCACCGGCAACCCAAAGCCACGGACATC | 3762369 | 3762428 |
| spoT_35 | TGCCCATTAAAGGTGCCGATGGCGTGCTGATCACCTTTGCGAAATGCTGCCGCCCTATTC | 3762429 | 3762488 |
| spoT_36 | CTGGCGACCCGATTATCGCCCACGTCAGCCCCGGTAAAGGTCTGGTGATCCACCATGAAT | 3762489 | 3762548 |
| spoT_37 | CCTGCCGTAATATCCGTGGCTACCAGAAAGAGCCAGAGAAGTTTATGGCTGTGGAATGGG | 3762549 | 3762608 |
| spoT_38 | ATAAAGAGACGGCGCAGGAGTTCATCACCGAAATCAAGGTGGAGATGTTCAATCATCAGG | 3762609 | 3762668 |
| spoT_39 | GTGCGCTGGCAAACCTGACGGCGGCAATTAACACCACGACTTCGAATATTCAAAGTTTGA | 3762669 | 3762728 |
| spoT_40 | ATACGGAAGAGAAAGATGGTCGCGTCTACAGCGCCTTTATTCGTCTGACCGCTCGTGACC | 3762729 | 3762788 |
| spoT_41 | GTGTGCATCTGGCGAATATCATGCGCAAAATCCGCGTGATGCCAGACGTGATTAAAGTCA | 3762789 | 3762848 |
| spoT_42 | CCCGAAACCGAAATTAATGTTTTATGAACCCAACACGTTATGCACGCATCTGCGAAATGC | 3762849 | 3762908 |

| Name | Sequence | Start | End |
|---|---|---|---|
| hslU_1 | AAATGGGGCCTTTCAGCCCCATCAAACAATGATGAAAATGATTGAACGCGATTATAGGAT | 4099848 | 4099907 |
| hslU_2 | AAAACGGCTCAGATCTTCATCTGCCACCAACGCATCCAGATGTTTGCTCACATAATCTGC | 4099908 | 4099967 |
| hslU_3 | GTCAATAGTGATATTTTGACCGCTTAAATCGCTGGCGTCGTAGGAAATCTCTTCCATTAA | 4099968 | 4100027 |
| hslU_4 | ACGCTCCAGAACAGTGTGTAAACGACGAGCACCGATGTTTTCGGTAGATTCGTTCACCTG | 4100028 | 4100087 |
| hslU_5 | CCATGCCGCTTCCGCGATGCGTTTAATACCGGAGTCGGTAAACTCGATATTTACGCCTTC | 4100088 | 4100147 |
| hslU_6 | AGTCGCCATCAGTGCTTTGTACTGCACGGTGATAGAGGCATTCGGCTCGGTCAGAATACG | 4100148 | 4100207 |
| hslU_7 | CTCGAAGTCGCTGGTGGTCAGCGCCTGCAGTTCAACGCGGATTGGCAGACGACCTTGCAG | 4100208 | 4100267 |
| hslU_8 | TTCCGGGATCAGGTCAGACGGTTTCGCAATCTGGAACGCGCCAGAAGCGATAAACAGAAT | 4100268 | 4100327 |
| hslU_9 | GTGGTCAGTTTTGACCATCCCGTGTTTGGTGGAAACGGTGCAACCTTCTACCAGCGGCAG | 4100328 | 4100387 |
| hslU_10 | CAGGTCACGCTGAACGCCTTCACGAGAAACATCCGGACCGGAAGACTCGCCGCGCTTACA | 4100388 | 4100447 |
| hslU_11 | GATTTTGTCGATTTCGTCGATAAACACGATCCCGTGCTGCTCAACAGCGTCGATAGCGTC | 4100448 | 4100507 |
| hslU_12 | TTGCTTCAGCTCTTCCGGGTTCACCAGTTTCGCCGCTTCTTCTTCAATCAGCAGCTTCAT | 4100508 | 4100567 |
| hslU_13 | GGCGTCTTTGATTTTCAGCTTACGCGCTTTTTGCTTCTGGCCGCCCAGGTTCTGGAACAT | 4100568 | 4100627 |
| hslU_14 | GGACTGCAGCTGGCTGGTCATCTCTTCCATGCCCGGAGGAGCCATAATTTCAACGCCCAT | 4100628 | 4100687 |
| hslU_15 | CGGTGCTGCGGCAAGATCGATCTCGATTTCTTTGTCATCAAGCTGGCCTTCACGCAGTTT | 4100688 | 4100747 |
| hslU_16 | TTTGCGGAATGCCTGACGAGCAGCGGACGGTTCCTGCTGCTGTTCGGTCTGTCCCCAGTT | 4100748 | 4100807 |
| hslU_17 | GTTTTTAGCAGGTGGGATCAGCACGTCGAGAATACGTTCTTCTGCCAGTTCTTCAGCGCG | 4100808 | 4100867 |
| hslU_18 | ATAACGGTTTTTCTCGATAGCCTGGACGCGTACCATTTTCACGGCGGCATCGGTCAGATC | 4100868 | 4100927 |
| hslU_19 | GCGAATAATAGAATCCACTTCCTTACCGACGTAGCCCACTTCGGTGAATTTGGTCGCTTC | 4100928 | 4100987 |
| hslU_20 | AACTTTGATGAACGGCGCATTCGCCAGCTTAGCCAGACGACGGGCGATTTCAGTTTTACC | 4100988 | 4101047 |
| hslU_21 | GACACCGGTCGGGCCGATCATCAGGATATTTTTCGGGGTCACTTCATGGCGCAGCTCTTC | 4101048 | 4101107 |
| hslU_22 | GTTGAGCTGCATGCGACGCCAGCGGTTACGCAGAGCAATCGCCACAGAACGCTTGGCGTT | 4101108 | 4101167 |
| hslU_23 | GTCCTGGCCGATGATGTGCTTATCCAGTTCGCTGACGATTTCGCGTGGGGTCATTTCAGA | 4101168 | 4101227 |
| hslU_24 | CATGGGAGATCCTTACGCTTTGTAGCTTAATTCTTCGATGGTGTGGAAATGGTTGGTATA | 4101228 | 4101287 |
| hslU_25 | GATGCAAATGTCGCCTGCAATATCCAACGCCTTTTCAGCAATTTCACGGGCGCTAAGTTC | 4101288 | 4101347 |
| hslU_26 | AGTGTTTTCTAACAGCGCGCGCGCCGCAGCCTGGGCGTAAGGGCCGCCGGAGCCGATAGC | 4101348 | 4101407 |
| hslU_27 | AATAAGATCGTTTTCTGGCTGCACCACGTCACCGTTACCGGTGATGATAAGCGATGCAGT | 4101408 | 4101467 |
| hslU_28 | TTCATCCGCGACTGCCAGCAGTGCTTCAAGTTTGCGCAGCATGCGATCGGTACGCCAGTC | 4101468 | 4101527 |
| hslU_29 | TTTTGCCAGCTCAACGGCGGCTTTGACCAGATGGCCCTGATGCATTTCCAGTTTACGTTC | 4101528 | 4101587 |
| hslU_30 | AAACAGTTCGAACAGCGTAAAAGCATCCGCAGTACCGCCCGCAAAGCCCGCGATGACTTT | 4101588 | 4101647 |
| hslU_31 | GTCGTTGTACAGACGGCGGACCTTTTTCACGTTGCCTTTCATTACGGTATTGCCCAACGT | 4101648 | 4101707 |
| hslU_32 | GGCCTGACCATCACCAGCGATGACCACATGGCCGTTACGGCGTACGCTTACTATAGTTGT | 4101708 | 4101767 |
| hslU_33 | CACGAGCTGACCCCTTGGTTACGAATACAGAGTACAAACCCCGTACAAAAGTACGGGGCA | 4101768 | 4101827 |
| hslU_34 | TAATGCAATTATAGATGGGGGGGATTTTGAGGGTTTCAACCCCCGGCGGCGAGCCGAATG | 4101828 | 4101887 |
| fabR_1 | GCCGGGGCCGGAACCTATTACTATGGCATCGTAATCGTAGGAATGTGGCATGGTAGGGCT | 4140601 | 4140660 |
| fabR_2 | TACCTGTTCTTATACATAAAAGCAACAGAATGGTAACATTTTATCGCGGGTAAGCCAATT | 4140661 | 4140720 |
| fabR_3 | GATCCCCGTCATTTATCTGGCTATATCCTGAGCGGCCTTTGCTTTGTCTGTTTCTTACTT | 4140721 | 4140780 |
| fabR_4 | TTGCCCTGACGTTTTATTGGATTTTTATCGACGATACTCTCCGTTTAAGCGGCAGGTTTC | 4140781 | 4140840 |
| fabR_5 | CGCTGTACGTAAAAGAACCGGCCAAAGAATTGCAGTAAATATGTTTTATTGCGTTACCGT | 4140841 | 4140900 |
| fabR_6 | TCATTCACAATACTGGAGCAATCCAGTATGTTCATTCTCTGGTATAGTGCCAGCAGTACT | 4140901 | 4140960 |
| fabR_7 | TTTGGCAAGGATTCAGACATCGTGATGGGCGTAAGAGCGCAACAAAAAGAAAAAACCCGC | 4140961 | 4141020 |
| fabR_8 | CGTTCGCTGGTGGAAGCCGCATTTAGCCAATTAAGTGCTGAACGCAGCTTCGCCAGCCTG | 4141021 | 4141080 |
| fabR_9 | AGTTTGCGTGAAGTGGCGCGTGAAGCGGGCATTGCTCCCACCTCTTTTTATCGGCATTTC | 4141081 | 4141140 |
| fabR_10 | CGCGACGTAGACGAACTGGGTCTGACCATGGTTGATGAGAGCGGTTTAATGCTACGCCAA | 4141141 | 4141200 |
| fabR_11 | CTCATGCGCCAGGCGCGTCAGCGTATCGCCAAAGGCGGGAGTGTGATCCGCACCTCGGTC | 4141201 | 4141260 |
| fabR_12 | TCCACATTTATGGAGTTCATCGGTAATAATCCTAACGCCTTCCGGTTATTATTGCGGGAA | 4141261 | 4141320 |
| fabR_13 | CGCTCCGGCACCTCCGCTGCGTTTCGTGCCGCCGTTGCGCGTGAAATTCAGCACTTCATT | 4141321 | 4141380 |
| fabR_14 | GCGGAACTTGCGGACTATCTGGAACTCGAAAACCATATGCCGCGTGCGTTTACTGAAGCG | 4141381 | 4141440 |
| fabR_15 | CAAGCCGAAGCAATGGTGACAATTGTCTTCAGTGCGGGTGCCGAGGCGTTGGACGTCGGC | 4141441 | 4141500 |
| fabR_16 | GTCGAACAACGTCGGCAATTAGAAGAGCGACTGGTACTGCAACTGCGAATGATTTCGAAA | 4141501 | 4141560 |
| fabR_17 | GGGGCTTATTACTGGTATCGCCGTGAACAAGAGAAAACCGCAATTATTCCGGGAAATGTG | 4141561 | 4141620 |

| Name | Sequence | Start | End |
|---|---|---|---|
| fabR_18 | AAGGACGAGTAATGAAACAAGCAAATCAAGATAGAGGTACGCTGCTGCTGGCGTTAGTTG | 4141621 | 4141680 |
| iclR_1 | TCTATTGCCACTCAGGTATGATGGGCAGAATATTGCCTCTGCCCGCCAGAAAAAGTCAGC | 4201680 | 4201739 |
| iclR_2 | GCATTCCACCGTACGCCAGCGTCACTTCCTTCGCCGCTTTAATCACCATTGCGCCAAACT | 4201740 | 4201799 |
| iclR_3 | CGGTCACGCGGTCATCGGTAATACGTGAAATCGGTCCGGAAATAGAAATTGCGGCAAACG | 4201800 | 4201859 |
| iclR_4 | GTTCACGGTGCTCATCGAAAATACACGCTGCAAGGCAACGTAGCCCCAGCGCATGTTCCT | 4201860 | 4201919 |
| iclR_5 | CATCGTCAAATGAATAACCCCGTTTGCGCGTTTGGGCGAGATCTTCTTTTAAATGCACAG | 4201920 | 4201979 |
| iclR_6 | GAGACACCAGCGTTGCGTGGGTATAGGCATGTAACCCTTTGCGGTGCAGCAGCTTCGTCA | 4201980 | 4202039 |
| iclR_7 | CCTGTTCTTCGCTCAGTTGGGCTAAAAAGGCTTTACCTGCACCGGAAGCGTGCATCGGCA | 4202040 | 4202099 |
| iclR_8 | ATTTACCGCCGATAGGCGCGGACATTCGCATCAGATGCGTACACTGTACCTGGTCGATAA | 4202100 | 4202159 |
| iclR_9 | TAATCGCTTCGTGATCGCTTTGATCAAGCACCGCCATATTGACCGTTTCGCCAGACTCTT | 4202160 | 4202219 |
| iclR_10 | CCATTAAATTGCGCAGGATAGGGTGAACAATCGCTAACAAATTACGGCTCTGGAGAAAGC | 4202220 | 4202279 |
| iclR_11 | TGCTGCCGACCATAAAGGCATGTGCGCCGATTGCCCAATGTCCCAGTTCGCCGACCTGAC | 4202280 | 4202339 |
| iclR_12 | GCACGAAACCCTGCTGTTGCATTGTGGTTAGCAGGCGGTGGGTCGTGGAATTGGGTAACC | 4202340 | 4202399 |
| iclR_13 | CGGCTTGTTGCGCCAGTTCCGTGAGTGCCACACTGCCATTGGATTCGGCAATCCACTCCA | 4202400 | 4202459 |
| iclR_14 | GTAATTTCAGGCCACGCGTTAAAGACTGAACCTGTCCAGTCGCTGGTGCGGTGGCAACGG | 4202460 | 4202519 |
| iclR_15 | CGGGTTTTCTGCCGCGTTTCGCGGGAATGGGTGCGACCATGACAGTCTCCTTTTTCTGTA | 4202520 | 4202579 |
| iclR_16 | TCGTGGAAATCATTTTCATTTTTATTGTTAGCTAATGCAATAGTTGCTGAACTGATCCGA | 4202580 | 4202639 |
| iclR_17 | TGAGTTAATGTTGAACAAATCTCATGTTGCGTGGTGGTCGCTTTTACCACAGATGCGTTT | 4202640 | 4202699 |
| nadR_1 | CTTAGCGTGTTCGACGACTTATAATGAGGAATACGGAGGGAGATATGTCGTCATTTGATT | 4615485 | 4615544 |
| nadR_2 | ACCTGAAAACTGCCATCAAGCAACAGGGCTGCACGCTACAGCAGGTGGCTGATGCCAGCG | 4615545 | 4615604 |
| nadR_3 | GTATGACCAAAGGGTATTTAAGCCAGTTACTGAATGCCAAAATCAAAAGCCCCAGCGCGC | 4615605 | 4615664 |
| nadR_4 | AAAAGCTGGAGGCGTTGCACCGTTTTTTGGGGCTTGAGTTTCCCCGGCAGAAGAAAACCA | 4615665 | 4615724 |
| nadR_5 | TTGGTGTGGTATTCGGTAAGTTCTACCCGCTGCATACCGGACATATCTACCTTATCCAGC | 4615725 | 4615784 |
| nadR_6 | GCGCCTGTAGCCAGGTTGACGAACTGCATATCATTATGGGTTTTGACGATACCCGCGATC | 4615785 | 4615844 |
| nadR_7 | GCGCGTTGTTTGAAGACAGCGCTATGTCGCAGCAGCCCACCGTGCCGGATCGTCTGCGCT | 4615845 | 4615904 |
| nadR_8 | GGTTATTACAAACTTTTAAATATCAGAAAAATATTCGCATTCATGCTTTCAACGAAGAGG | 4615905 | 4615964 |
| nadR_9 | GCATGGAGCCGTATCCGCACGGCTGGGATGTGTGGAGCAACGGCATCAAAAAGTTTATGG | 4615965 | 4616024 |
| nadR_10 | CTGAAAAGGGGATTCAGCCGGACCTGATCTACACCTCGGAAGAAGCCGATGCGCCACAGT | 4616025 | 4616084 |
| nadR_11 | ATATGGAACATCTGGGGATCGATACGGTGCTGGTCGATCCGAAACGTACCTTTATGAGTA | 4616085 | 4616144 |
| nadR_12 | TCAGCGGTGCGCAGATCCGCGAAAACCCGTTCCGCTACTGGGAATATATTCCTACGGAAG | 4616145 | 4616204 |
| nadR_13 | TGAAGCCGTTCTTTGTACGTACCGTGGCGATCCTTGGTGGCGAGTCGAACGGTAAATCCA | 4616205 | 4616264 |
| nadR_14 | CCCTGGTAAACAAACTTGCCAATATCTTCAATACCACCAGTGCGTGGGAATATGGTCGCG | 4616265 | 4616324 |
| nadR_15 | ATTATGTCTTTTCACACCTCGGCGGTGATGAGATCGCATTGCAGTATTCCGATTACGATA | 4616325 | 4616384 |
| nadR_16 | AAATCGCGCTGGGCCACGCACAATACATTGATTTTGCAGTGAAATATGCCAATAAAGTGG | 4616385 | 4616444 |
| nadR_17 | CGTTTATTGACACTGATTTTGTCACTACCCAGGCGTTCTGCAAAAAGTACGAAGGGCGTG | 4616445 | 4616504 |
| nadR_18 | AGCATCCGTTCGTACAGGCGTTGATTGATGAATACCGTTTCGATCTGGTGATCCTGCTGG | 4616505 | 4616564 |
| nadR_19 | AGAACAACACGCCGTGGGTGGCGGATGGTTTACGCAGCCTCGGCAGTTCGGTGGATCGCA | 4616565 | 4616624 |
| nadR_20 | AAGAGTTCCAGAACTTGCTGGTGGAGATGCTGGAAGAGAACAATATCGAATTCGTGCGGG | 4616625 | 4616684 |
| nadR_21 | TTGAAGAGGACGATTATGACAGCCGTTTCCTGCGCTGCGTGGAGCTGGTGCGGGAGATGA | 4616685 | 4616744 |
| nadR_22 | TGGGGGAGCAGAGATAACCGCGATGAAACGGCTCAAAGGCGAGGTATAAAATAAGTTTTT | 4616745 | 4616804 |

All sequences were ordered as xGen Lockdown probes with a 5′ biotin tag from Integrated DNA Technologies. Start and end are the positions where the first and last base map to in the *E. coli* REL606 genome (GenBank: NC_012967.1).

**Table S3.** Blocker sequences

| Name | Sequence |
|---|---|
| Universal Blocker | AATGATACGGCGACCACCGAGATCTACACTCTTTCCCTACACGACGCTCTTCCGATCT |
| Degenerate Sample Barcode Blocker | CAAGCAGAAGACGGCATACGAGAT**NNNNNN**GTGACTGGAGTTCAGACGTGTGCTCTTCCGATCT |

**Red bases** anneal to the sample barcodes. They were synthesized with a mixture of all four bases so that one blocking oligo could be used in pulldowns in which multiple different sample barcodes were present.
